# Dynamic changes to the plastoglobule lipidome and proteome in water-deficient maize

**DOI:** 10.1101/2025.05.26.656201

**Authors:** Elsinraju Devadasu, Anthony L. Schilmiller, Natali Gonzalez, Peter K. Lundquist

## Abstract

Drought represents one of the most severe challenges faced by agriculture and leveraging resources to promote crop resilience is critical. The plastoglobule lipid droplets of chloroplasts, present in all photosynthetic organisms, are suggested to be a major orchestrator of adaptive responses to environmental perturbations, thus representing a potentially significant, untapped target for enhancement of crop resilience. Yet, the functions of plastoglobules are unclear and their molecular composition incompletely described. Here, we provide a thorough investigation of the protein and lipid compositions of plastoglobules and thylakoids at six time-points over the course of a water-deficit and recovery treatment in B73 inbred maize. Our results establish the prominent components of the plastoglobule polar lipid surface and neutral lipid interior in an important crop species, including the presence of mono- and di-galactosyl diacylglycerol lipids enriched in saturated acyl groups, and the prevalence of various triacylglycerols and plastoquinone-9 derivatives. Quantitative proteomics identifies prominent Fibrillins and Activity of bc_1_ Complex Kinases at the plastoglobule as well as many proteins with known or putative roles in prenyl-lipid and redox metabolism. A remarkably high proportion of the Fibrillin 4 on the plastoglobules coincided with a preponderance of plastoquinone-9, supporting a role for Fibrillin 4 in plastoquinone accumulation at plastoglobules. Collectively, our results provide a solid foundation for the study of plastoglobules in crop plants.

## Introduction

Drought is a major threat to agriculture that is predicted to increase in frequency and severity (Agency; Center; Alexander et al., 2006; Sillmann et al., 2013). The resultant water deficit inflicted within plant tissues imposes severe limitations on crop productivity through negative impacts on numerous plant processes such as growth, development and photosynthesis (Seleiman et al., 2021; Sato et al., 2024). The impacts of drought on plant physiology are a manifestation of a complex interplay of molecular and cellular responses. The phytohormone abscisic acid (ABA) is a major regulator of drought responses and induces stomatal closure upon perception of low water potential within the leaf. Phospholipase Dα1 is a mediator of the ABA-dependent stomatal closure and, at later stages of drought, is also associated with membrane degradation (Hong et al., 2008). Indeed, large-scale membrane remodeling is a hallmark of water-deficit responses in plants and major ultrastructural changes within the leaf coincide with the progressive reduction of relative water content (RWC). These membrane remodeling activities can include autophagy and accumulation of triacylglycerols (TAGs) within lipid droplets (Munne-Bosch and Alegre, 2004; Perlikowski et al., 2016; Ferreira et al., 2021; Hickey et al., 2022).

Much of a plant’s water-deficit responses are centered in the chloroplast, which has emerged as a primary site of abiotic stress perception and an important component of stress adaptation and tolerance (Estavillo et al., 2011; Woodson and Chory, 2012; Chan et al., 2016; de Souza et al., 2017; Grubler et al., 2021; Kachroo et al., 2021; Liebers et al., 2022). ABA biosynthesis is initiated within the chloroplast using photosynthetic carotenoid pigments as precursor substrate. The stomatal closure induced by ABA will also lead to a reduction in C assimilation rates requiring strong reduction in light harvesting and electron transport to balance provision of chemical energy and reducing power with its use in the Calvin-Benson-Bassham cycle.

The down-shift in light harvesting is accomplished through multiple mechanisms. For example, a strong induction of the xanthophyll cycle, an important component of the energy-dependent (qE) form of non-photochemical quenching (NPQ) allows dissipation of excess light energy that cannot be used productively in electron transport (Muller et al., 2001; North et al., 2005). Additionally, major impacts on the photosynthetic machinery can be seen, including disassembly of Photosystem II (PSII) supercomplexes and, at more severe levels of drought, disassembly of the PSI supercomplexes (Hu et al., 2023). The impacts on photosystem assembly coincide with greater levels of PSII photoinhibition and development of excess (*i.e.*, uncontrolled) reactive oxygen species (ROS) generation and lipid peroxidation (Munne-Bosch and Cela, 2006; Bertamini et al., 2007; Wang et al., 2018; Nosalewicz et al., 2022). The ratio of monogalactosyl diacylglycerol (MGDG) and digalactosyl diacylglycerol (DGDG) within the thylakoid membrane can also change, as shown in two maize (*Zea mays*) cultivars which increased their DGDG/MGDG ratios under drought (Chen et al., 2018). Corresponding to these broad impacts on chloroplast biochemistry, substantial effects on chloroplast ultrastructure are also observed. Under severe drought stress, chloroplasts appear swollen with reduced envelope membrane integrity, thylakoid membranes unstack resulting in fewer or smaller grana, and plastoglobule lipid droplets proliferate (Grigorova et al., 2012; Carrera et al., 2021; Chen et al., 2022; Aliyeva et al., 2023).

Plastoglobules have emerged as likely regulatory hubs for chloroplast metabolism and signaling during stress and, as such, may present attractive targets for the enhancement of crop resilience (Lundquist et al., 2020). Plastoglobules are present in the chloroplasts of all photosynthetic organisms and are formed by a blebbing of the outer leaflet of the thylakoid membrane, such that the periphery of each plastoglobule is a single, monolayer membrane leaflet (Austin et al., 2006). Thus, the plastoglobule comprises a purely hydrophobic interior accommodating non-polar lipid species, while amphipathic lipids and all proteins are associated with the plastoglobule’s periphery. Despite the physical association with the thylakoid membrane, the plastoglobule comprises a distinct lipid and protein composition that is rich in prenyl-lipid compounds (*e.g.,* carotenoids, tocochromanols and quinones), various neutral lipids such as fatty acid phytyl esters and xanthophyll esters, and a protein-studded periphery.

The dynamic morphology of plastoglobules and their ubiquitous nature throughout the plant kingdom suggests they hold critical role(s) in stress adaptation. Experimental studies of *A. thaliana* mutant lines disrupted in genes whose products localize to the plastoglobule have supported this assumption. The function(s) of the plastoglobule remain largely enigmatic but are likely driven by the functions of the associated proteins. A quantitative “core” proteome of plastoglobules isolated from *Arabidopsis thaliana* has been described using mass-spectrometry-based proteomics, identifying 32 proteins that were highly-enriched (or exclusively localized) to the plastoglobule (Lundquist et al., 2012). This included seven members of the Fibrillin (FBN) protein family characterized by a lipid-binding Lipocalin domain, six Activity of bc_1_ protein kinases (ABC1Ks), and numerous other characterized or putative metabolic enzymes related to redox regulation and prenyl-lipid metabolism. Comparative analysis of plastoglobules from control and high-light stressed *A. thaliana* plants revealed tailored changes to the proteome and prenyl-lipidome consistent with mediation of senescent-like processes, including the generation of the phytohormone, jasmonic acid (JA) and remobilization of carotenoids from the photosynthetic machinery to the plastoglobules (Espinoza-Corral et al., 2021). Moreover, investigation of plastoglobules from an *A. thaliana* double mutant disrupting two plastoglobule-localized ABC1Ks revealed a remobilization of the plastid-localized enzymes of JA biosynthesis to plastoglobules (Lundquist et al., 2013). These results highlight the dynamic nature of plastoglobules under stress and point to their role as a platform for metabolism and signaling, consistent with their proliferation under diverse stresses.

Despite the advancements in our understanding of plastoglobules outlined above, the investigation of control and light-stressed plastoglobules from *A. thaliana* described above represents the only direct comparative analysis of stressed and unstressed plastoglobule proteomes and lipidomes in the literature. Furthermore, the characterization of the plastoglobule composition has not previously been extended to a crop plant and, beyond the composition of prenyl-lipids, their lipidome is still poorly understood. A detailed characterization of the plastoglobule composition and how it changes in response to stresses would provide valuable insights into the adaptation and functions of the plastoglobule under stress. In this work, we ask what are the lipid and protein components of the *Z. mays* plastoglobules, how does this compare to the changes at the adjacent thylakoid membrane, and what accounts for the substantial swelling in size and abundance seen under stress.

In this study, we implemented a water-deficit and recovery time course treatment with three-week-old B73 inbred maize and investigated the ultrastructural behavior and molecular composition of plastoglobules and thylakoids at six different time points. Our results offer a detailed view of the molecular composition of plastoglobules that includes their surface polar lipid and interior neutral lipid composition, revealing targeted, rather than whole-sale, changes to the plastoglobule likely related to membrane remodeling. We further demonstrate that derivatives of plastoquinone (PQ-9) comprising hydroxyl or acyl groups on the prenyl tail and polar head are prominent components of the plastoglobule, whose functions remain to be determined. Our efforts will pave the way to rationally employ plastoglobules for better stress tolerance capacity in plants.

## RESULTS

### Optimization of the water deficit treatment of B73 inbred maize

A water-deficit treatment was imposed on B73 inbred maize three weeks after germination by withholding water. To establish a proper time course for our experimentation, we carried out pilot experiments monitoring the leaf relative water content (RWC; %) and photosynthetic traits. In particular, we sought a length of time that is sufficient to induce a clear stress to the plant that imposes a negative impact on photosynthetic capacity, but from which the plant could recover upon re-provision of water. We also used larger pots (9 cm diameter and 10 cm depth) containing only one individual maize plant in each pot to minimize pleiotropic effects from overly dense root growth or rapid water drainage. With this experimental setup we found that the leaf RWC of mature (*i.e.* collared) leaves was maintained around 97-100% for over two weeks without provision of water but dropped sharply between 16 and 17 days to *ca.* 65% leaf RWC (**Figure 1B, Supp Table S1**). This coincided with the emergence of chlorosis, leaf wilting, and curling at leaf tips (**Figure 1A, Supp Fig S1**), although chlorophyll and carotenoid levels were maintained at a whole leaf level when measuring on a leaf fresh weight basis (**Supp Table S2**). In addition, a decline in photosynthetic performance, including depressed Phi2 and F_v_^’^/F_m_^’^ values indicative of photoinhibition at PSII (Maxwell and Johnson, 2000; Baker, 2008) as well as elevated NPQ_t_ values (Tietz et al., 2017), were seen at 16 and 17 days (**Figure 1C & D, Supp Table S3**). Consistent with the elevated NPQ_t_, ratiometric quantification of immunoblots indicated elevated levels of PsbS, a component of the NPQ response in higher plants (**Figure 1E & F**) (Li et al., 2000; Teardo et al., 2007; Kiss et al., 2008). Re-provision of water at the 17 dWW stage led to a rapid (*i.e.* within one day) recovery of leaf RWC and restoration of photosynthetic parameters and PsbS levels (**Figure 1, Supp Tables S1 & S3**). In contrast, prolonging the water deprivation for an additional day saw a further steep decline in leaf RWC and onset of necrotic leaf tissue (data not shown), at which point provision of water could not restore photosynthetic performance or plant survival. The results of our pilot experiments indicated that withholding water from three-week-old maize for 17 days imposed a stress to the plants that was sufficiently severe to impact photosynthetic performance and induce leaf chlorosis at tip margins but maintain the capacity for full physiological recovery upon re- provision of water. Thus, this experimental time course provided a robust design for us to test the role of maize plastoglobules in the plant’s adaptation to water-deficit stress and subsequent recovery. We selected six time points across the stress and recovery phases of the time course for detailed investigation of cellular ultrastructure and protein and lipid compositions of isolated plastoglobules and thylakoids: 7, 12, 16, and 17 days without water (dWW), and 1 and 3 days after re-watering (dRW). We elected to start our analyses with the 7 dWW time point because the well-watered control maize is watered in 7-day intervals and no stress is anticipated within this interval, supported by our leaf RWC and photosynthetic measurements (**Figure 1, Supp Fig S1, Supp Tables S1 and S3**). Thus, we consider the 7 dWW time point as the un-stressed reference point.

**Figure 1.**
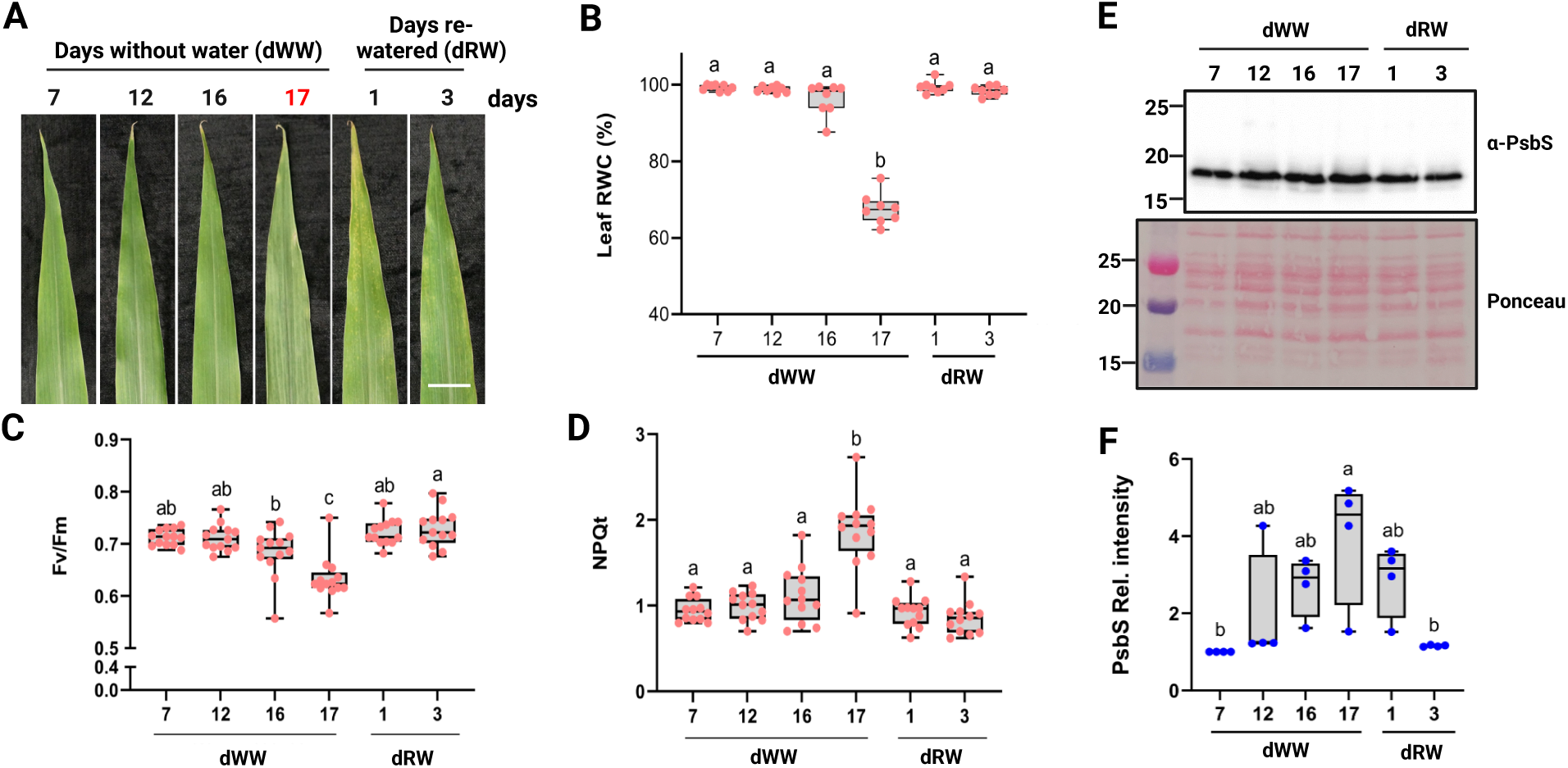
Phenotypic and photosynthetic measurements of maize leaves during the water deprivation and recovery time course. **A)** Representative photographs of the fourth leaf of maize plants at specified time points during the time course (scale bar = 5 cm). The peak stress time point, 17 days without water (17 dWW) is highlighted with red font. **B)** Leaf relative water content (RWC) of the fourth leaf at specified time points. n = 8. **C-D)** Photosynthetic traits measured with a handheld MultiSpeQ chlorophyll fluorometer. n = 12 **E)** Immunoblot of total leaf protein extracts using anti-PsbS antiserum. PsbS is a stress-inducible protein and regulator of energy-dependent non-photochemical quenching (NPQ). **F)** Densitometric measurements of the immunoblot. n = 4. Statistical significance of differences was assessed using one-way analysis of variance (ANOVA) with p < 0.001, indicated by lowercase letters.

### Manifestation of stress in the leaf proteome

Before preforming our time-course investigation of plastoglobules and thylakoids we developed a broad-level view of water-deficit effects by determining the quantitative total leaf proteome at the reference unstressed time-point (7 dWW) and the peak water-deficit stress time-point (17 dWW) (**Figure 2, Supp Table S4**). Protein was extracted from total above-ground leaf tissue, in-gel digested and analyzed with a Q-Exactive HF mass spectrometer connected in-line with nano-liquid chromatography (nanoLC-MS/MS). MS/MS raw files were searched against the *Zea mays* B73 reference genome version 3 using the Andromeda search engine within the MaxQuant software (Cox and Mann, 2008; Cox et al., 2011) with peptide- and protein-level false discovery rates of 1% using normalized label free quantification (nLFQ) (Cox et al., 2014). We identified a total of 3180 proteins or protein groups (hereafter, “proteins”) across all replicates and time points.

**Figure 2.**
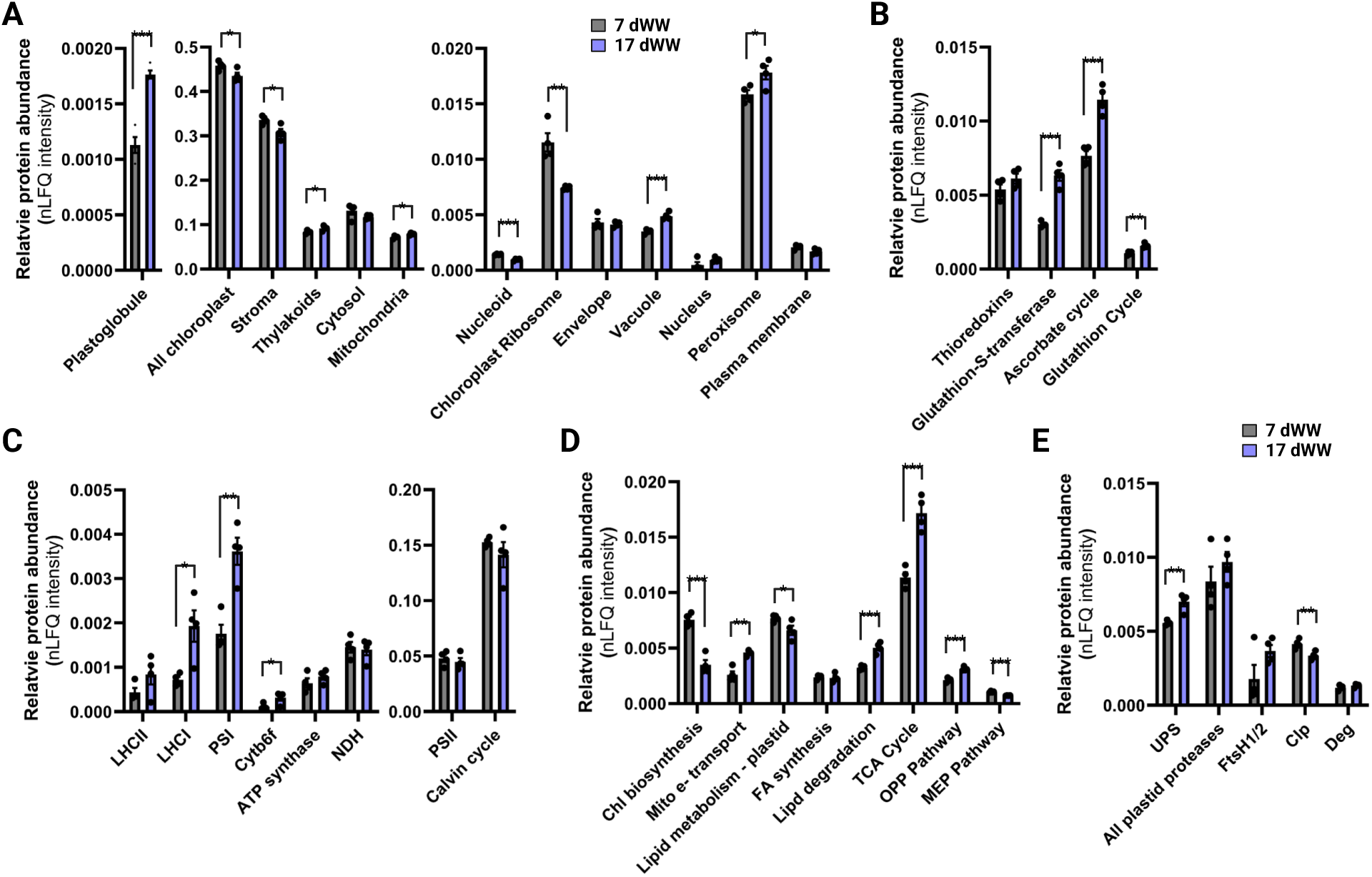
Comparison of the leaf proteomes under 7 and 17 days without water.

Evidence of stress could be clearly seen in the leaf proteome at 17 dWW. A substantial increase in oxidative stress machinery, including Glutathione-*S*-transferase proteins, and components of the ascorbate and glutathione cycles, was consistent with acute oxidative stress within the cell (**Figure 2B**). This coincided with an increase in the Ubiquitin Proteasome System, responsible for protein turnover within the cytosol, along with proteins involved in lipid degradation (**Figure 2D and E**). A heightened demand for energy was exhibited in the increased protein abundance of the mitochondrial electron transport chain, tricarboxylic acid (TCA) cycle and the oxidative pentose phosphate (OPP) pathway, possibly fueling energy-intensive adaptive responses such as the generation of compatible solutes and ROS-scavenging compounds.

Consistent with the increase in catabolic reactions, total protein of the peroxisomes and vacuole, compartments associated with catabolic processes, were also substantially elevated (**Figure 2A**). As expected, we also found a substantial increase in abundance of the plastoglobule proteome. More surprisingly, total thylakoid protein abundance increased under the stress (**Figure 2A**), despite the clear down-shift in chlorophyll biosynthesis enzymes (**Figure 2D**). This increase appeared to be driven predominantly by components of the Photosystem I (PSI) and Light-harvesting complex I (LHCI) complexes, and to a lesser extent the Cytochrome b_6_f (Cyt b_6_f) complex (**Figure 2C**). The levels of the other photosynthetic complexes (including the Calvin-Benson cycle) remained essentially flat. Despite these trends, the overall protein abundance of the chloroplasts declined, manifested primarily at the stroma, as well as the nucleoids and the ribosomes (**Figure 2A**). Collectively, the results of our quantitative leaf proteomics reveal evidence of acute oxidative stress as well as induction of various catabolic pathways and energy generation that is possibly driving adaptive responses to the water-deficit conditions.

### Effects of water-deficit on cellular ultrastructure

Swelling and/or proliferation of plastoglobules are a hallmark response to both biotic and abiotic stresses (van Wijk and Kessler, 2017; Zechmann, 2019; Lundquist et al., 2020). To determine how our water deprivation stress impacted plastoglobule abundance and morphology over our time course, and to observe other impacts on cellular ultrastructure, we collected transmission electron micrographs of chemically fixed leaf tissue of both mesophyll (M) and bundle sheath (BS) cells at each of the six time points (**Figure 3**). Chloroplast size decreased in both M and BS cells (**Table 1, Supp Fig S2, Supp Table S5**). At the initial time point, plastoglobule abundance was greater in M cells than BS cells (*ca.* 0.31 µm^-2^ chloroplast area versus 0.21 µm^-2^ chloroplast area, respectively). Strikingly, the plastoglobule abundance (number per um^2^) increased substantially in the M cells (but not BS cells) during the stress, reaching *ca.* 0.72 µm^-2^, and recovered only modestly upon re-watering. Plastoglobule size increased in both M and BS cells, increasing the average cross-sectional diameter from about 90 nm to 122 nm. Notably, this increase in size appears to continue even after re-watering (1 and 3 dRW). It can be seen that the size and abundance of plastoglobules in both cell types is lesser than what we have previously reported within five field-grown maize hybrids (Ying et al., 2023). The lower size and abundance likely reflect the younger developmental stage (V5-V6, *versus* V17 in Ying, *et al*.), as well as the inbred germplasm and chamber conditions. It is also notable that the size and abundance of plastoglobules in our study are quite comparable between the two cell types despite the functional division of the cells and distinct morphology of the chloroplasts (*e.g.* much more elongated chloroplasts with a general lack of thylakoid membrane stacking in BS cells). Together, our ultrastructural results demonstrate that plastoglobule morphology responds to the water deprivation by increasing plastoglobule abundance (preferentially in M cells) or increasing the plastoglobule size (in both M and BS cells), consistent with the increased plastoglobule protein abundance; trends which are generally not reversed within three days after re-watering.

**Figure 3.**
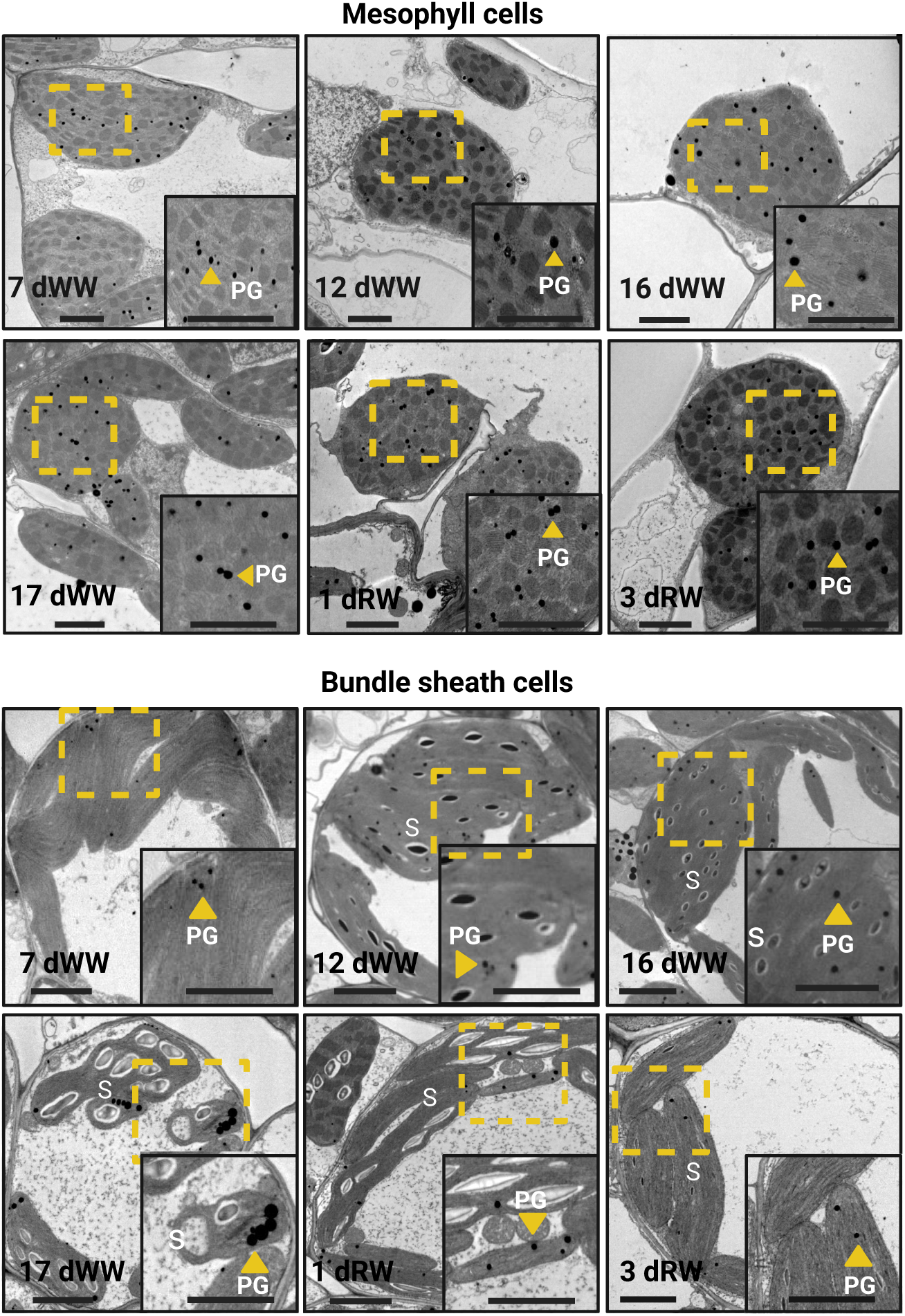
Cellular ultrastructure of mesophyll and bundle sheath cells during the water- deprivation-and-recovery time course. Representative transmission electron micrographs of mesophyll cells (top) and bundle sheath cells (bottom) at each of the six time points of the study. Scale bars of main micrographs are 2 μm and of insets are 500 nm. Letters highlight selected ultrastructural features, namely the plastoglobules (PGs) and starch grains (S).

**Table 1.**
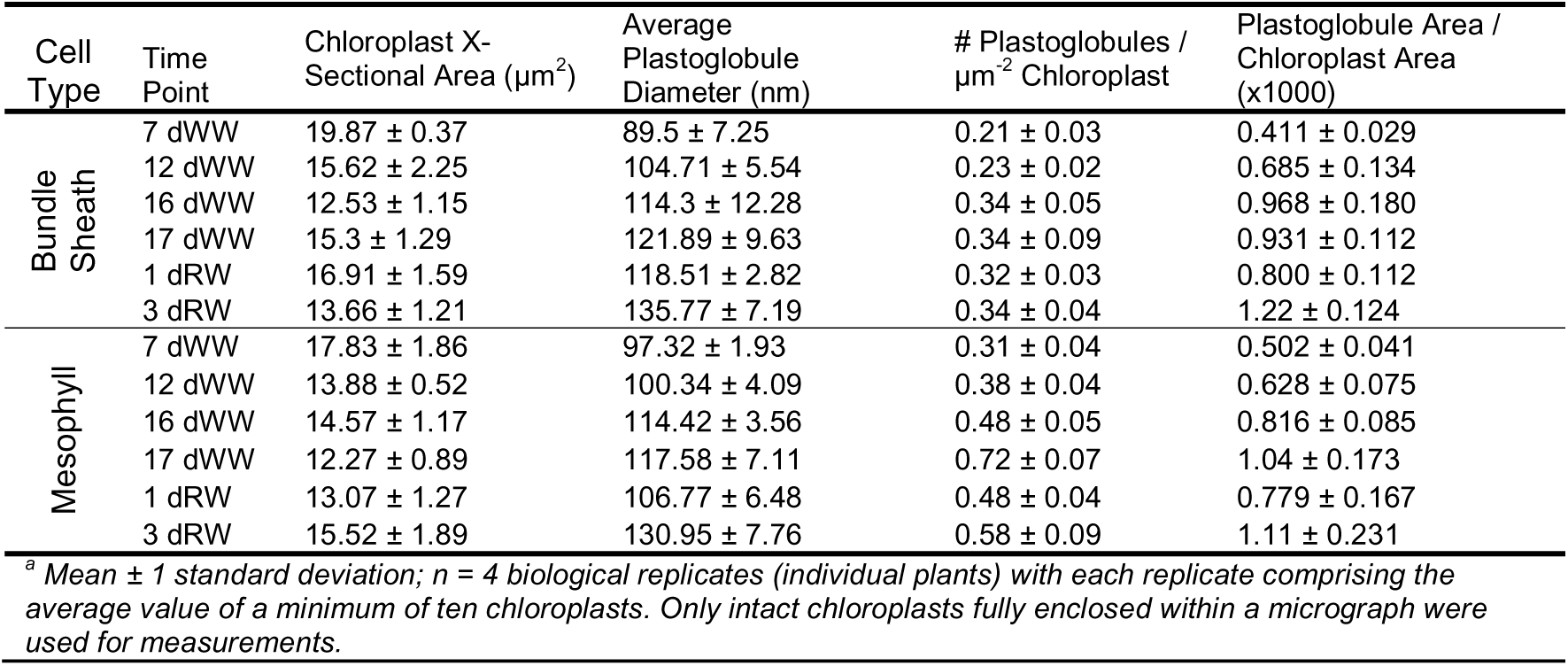
Ultrastructural measurements of *Z. mays* chloroplasts and plastoglobules during a water-deficit stress time-course^a^.

### Isolation of plastoglobules and thylakoids

Because we hypothesized that remodeling of the plastoglobule is tightly connected to corresponding effects at the thylakoid membrane, we wished to closely investigate the composition of thylakoids and plastoglobules in parallel. Accordingly, thylakoids and plastoglobules were isolated from the same leaf tissue at each of the six time points of the water deprivation treatment. Consistent with the proliferation of size and number of plastoglobules that we observed by TEM, the yield of isolated plastoglobule material increased substantially during the time course (**Supp Fig S3A**). Notably, we elected to analyze aliquots of thylakoid not subsequently sonicated (our method to separate plastoglobules from thylakoids prior to flotation of the plastoglobules) to avoid artefactual effects on the thylakoids, even though this means that the physically associated plastoglobules were not removed and were instead co-isolated with the thylakoids. We feel this will have only a trivial impact on our study of the thylakoid because of the immense quantity of thylakoid within the leaf tissue that dwarfs the contributions of plastoglobules. Immunoblotting of suitable marker proteins (PsbA [D1] for thylakoids and Fibrillin1a [FBN1a] for plastoglobules) indicated effective enrichment of both sub-compartments (**Supp Fig S3B**). Consistent with our observations of plastoglobule morphology, the levels of FBN1a protein associated with the thylakoid samples increased approximately two-fold over the time course of water deprivation (**Supp Fig S3B**). Isolated thylakoids and plastoglobules were subsequently used for detailed characterization, as described below.

### Quantitative proteomics point to tailored changes and dynamic remobilization at plastoglobules and thylakoids

Plastoglobule and thylakoid samples were processed and analyzed as described for the total leaf proteome samples above. We identified a total of 748 proteins or protein groups (hereafter, proteins) across all thylakoid samples of all time points (**Supp Table S6**) and 1079 proteins across all plastoglobule samples of all time points (**Supp Table S7**).

In the thylakoid samples, a total of 42 proteins changed in abundance between 7 dWW and 17 dWW with statistical significance (adjusted p-value < 0.05), 27 of which were then restored to pre-stress levels upon re-watering (**Supp Table S6**). Notably, seven of these proteins were plastoglobule proteins, including ABC1K9 (increased >5-fold), PG18 (increased > 4-fold) and FBN1a (increased nearly 4-fold). Consistent with the initiation of chlorophyll turnover, Pheophorbide a Oxygenase also increased at the thylakoid over 10-fold, while the chlorophyll biosynthesis proteins, Mg^2+^-Protoporphyrin IX Chelatase, subunits I and H, were lost from the thylakoid.

Looking at the impact of the stress on the major complexes of the electron transport chain and light harvesting within the thylakoid samples we found that, in general, quantitative changes mirrored what was seen at the total leaf level described above, albeit more muted. The PSI complex increased significantly during stress (*ca.* 31% increased from 7 dWW to 17 dWW) and partially recovered after re-watering (**Figure 4A, Supp Table S6**). In contrast, the ATP synthase complex and the NAD(P)H dehydrogenase (NDH) complex decreased during the stress, particularly at the 17 dWW time point and recovered fully by 1 dRW (**Supp Fig S4, and Supp Table S6**). We then looked within the thylakoid proteome for other indicators of stress response and noted that two members of the LHC-like (LIL) family, LIL3:1 and LIL3;2, were steadily reduced over the course of the stress, reaching values less than 50% of pre-stress by 17 dWW, and then partially recovered during re-watering (**Figure 4A**). Consistent with the general stability of the major thylakoid complexes, thylakoid proteases (including DegPs, FtsHs, and EGYs) were unchanged during the time-course. Thylakoid-localized Ascorbate peroxidases (tAPXs), as well as Glutaredoxins and Peroxidredoxins, were generally unchanged during the time course, although m-type and f-type Thioredoxins (Trxs) were seen to increase during the initial days of the stress, before dropping at peak stress (17 dWW) and again increasing during re-watering (**Supp Fig S4, and Supp Table S6**). It must be noted that, while these trends were observed with both Trx types, the changes were statistically significant only in the case of the m-type Trxs due to higher biological variability seen with the f-type Trxs. In sum, impacts of the stress can be seen within the thylakoid membrane proteome, particularly regarding impacts on the PSI, ATP synthase and NDH complexes, the LIL3 proteins and the Trxs, although broad changes to the photosynthetic machinery were not observed. This indicates that the plants have not triggered senescent-like processes but are initiating specific stress adaptive responses to the water deficit, consistent with the ability for the plants to recover rapidly upon re-watering.

**Figure 4.**
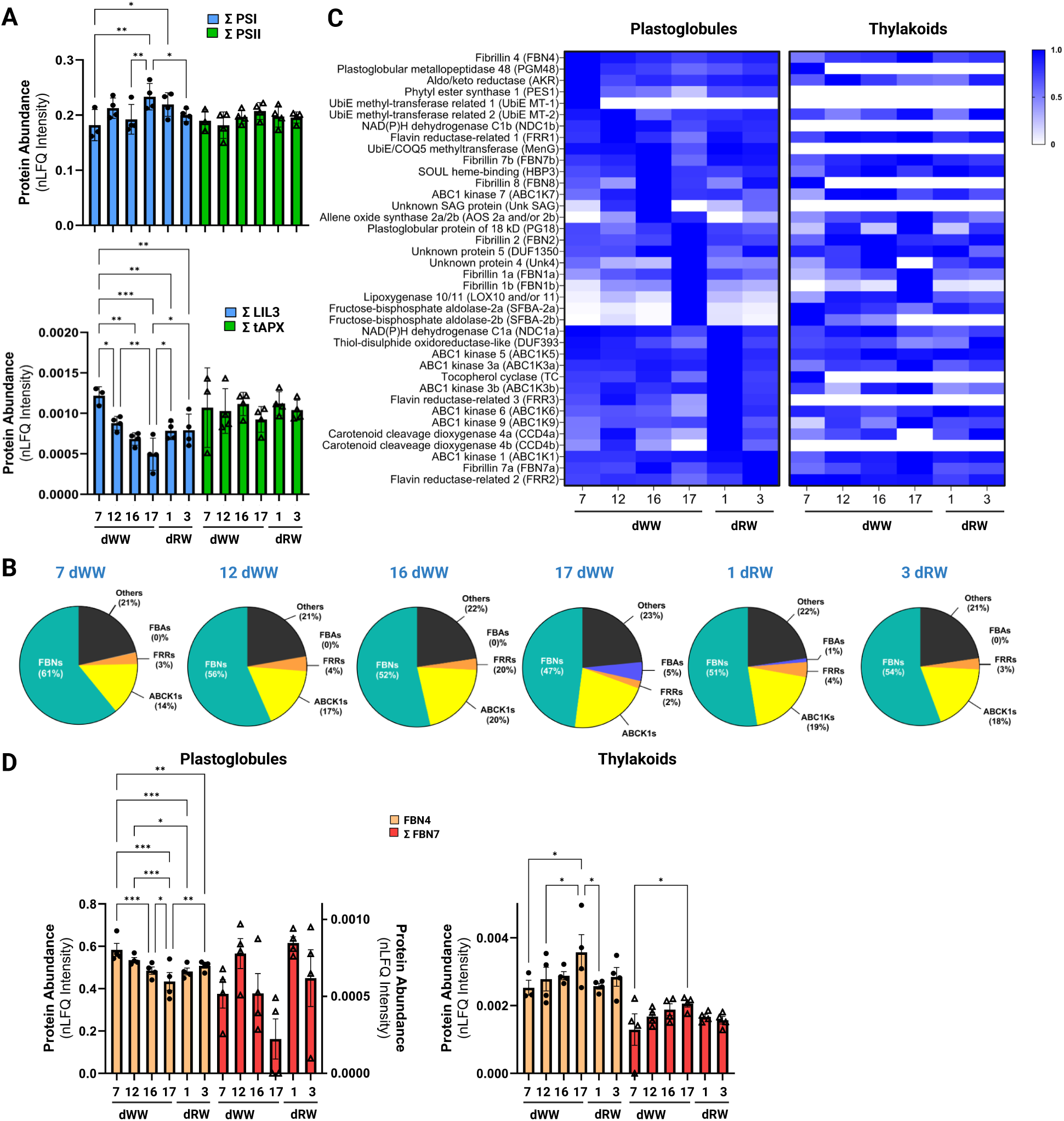
Quantitative proteomics of isolated Z. mays plastoglobules and thylakoids during the waterdeprivation-and-recovery time course. **A)** Summed nLFQ intensities of all detected subunits of specified photosynthetic complexes (Photosystem I and II [PSI and PSII] or protein families (LHCII-like proteins [LIL3] and thylakoid-associated Ascorbate Peroxidases [tAPXs]) in thylakoid samples. **B)** Relative proportions of the Z. mays plastoglobule proteome with selected protein families called out, namely the Fibrillin (FBN), Absence of bc1 Complex protein kinases (ABC1K), Fructose-bisphosphate Aldolase (FBA), and Flavin Reductase-related (FRR) protein families. **C)** Heatmap of normalized nLFQ intensities of all 38 ’core’ proteins of the Z. mays plastoglobule proteome. Each protein is normalized to its maximal abundance within each sub-compartment such that each protein will have its peak abundance of 1.0 at one of the six time points. Proteins are sorted by the time point at which they hold maximal protein abundance in the plastoglobule samples. **D)** nLFQ intensities of maize Fibrillin 4 (FBN4) and the sum of the Fibrillin 7 paralogs (Σ FBN7a + FBN7b) in plastoglobules (left) and thylakoids (right). Graphs plot mean ± 1 standard error of the mean, n = 3 (7 dWW) or 4 (all other time points) biological replicates (i.e., individual plants). ΣFBN7 in the plastoglobule sample is plotted on the secondary y-axis. Statistically significant differences within protein groups were determined in all panels using an ordinary two-way ANOVA; * p < 0.033, ** p < 0.002, *** p < 0.001.

The quantitative plastoglobule proteome of maize has not previously been described in the literature and the sensitivity of modern mass spectrometry-based proteomics means that even extremely low abundant contaminant proteins are readily identified in our plastoglobule isolations. Thus, to distinguish *bona fide* plastoglobule proteins from contaminant proteins in our plastoglobule isolations, we relied on the well-established *A. thaliana* plastoglobule proteome (Lundquist et al., 2012; Espinoza-Corral et al., 2021), as well as manually-curated plastoglobule localizations from the Plant Proteome Database (PPDB) (Sun et al., 2009). Clear homologs of each of the *A. thaliana* plastoglobule proteins are encoded in the maize genome, all but one of which were identified in our plastoglobule isolations (**Table 2, Supp Table S8**). The exception was the stress-induced Esterase/Lipase/Thioesterase 4 (ZmELT4; GRMZM5G817559), as noted below. Homology to *bona fide* plastoglobule proteins of *A. thaliana*, combined with their identification in isolated maize plastoglobules, led us to define a set of 38 proteins comprising the *Z. mays* plastoglobule proteome (**Table 2, Supp Table S8**).

**Table 2.**
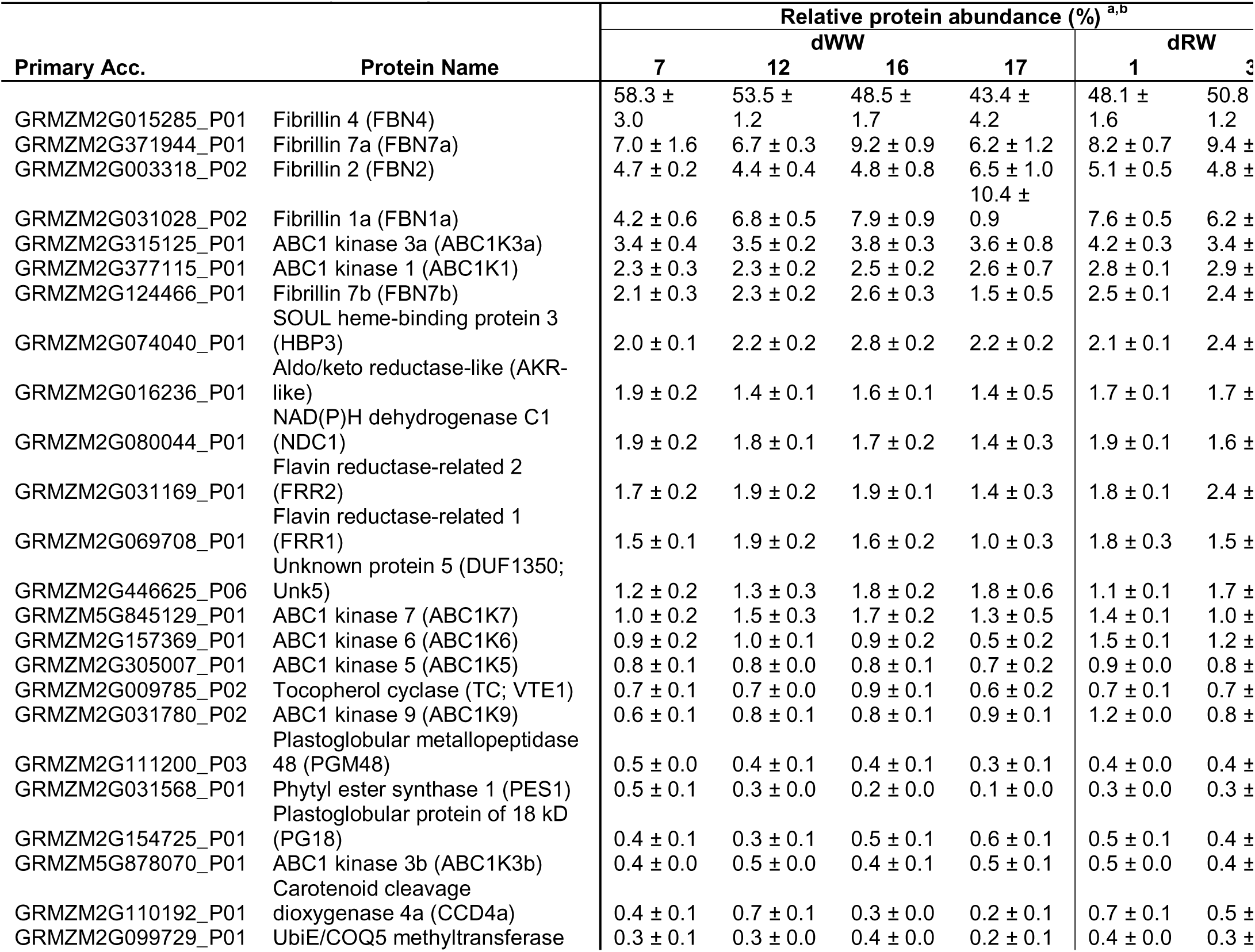
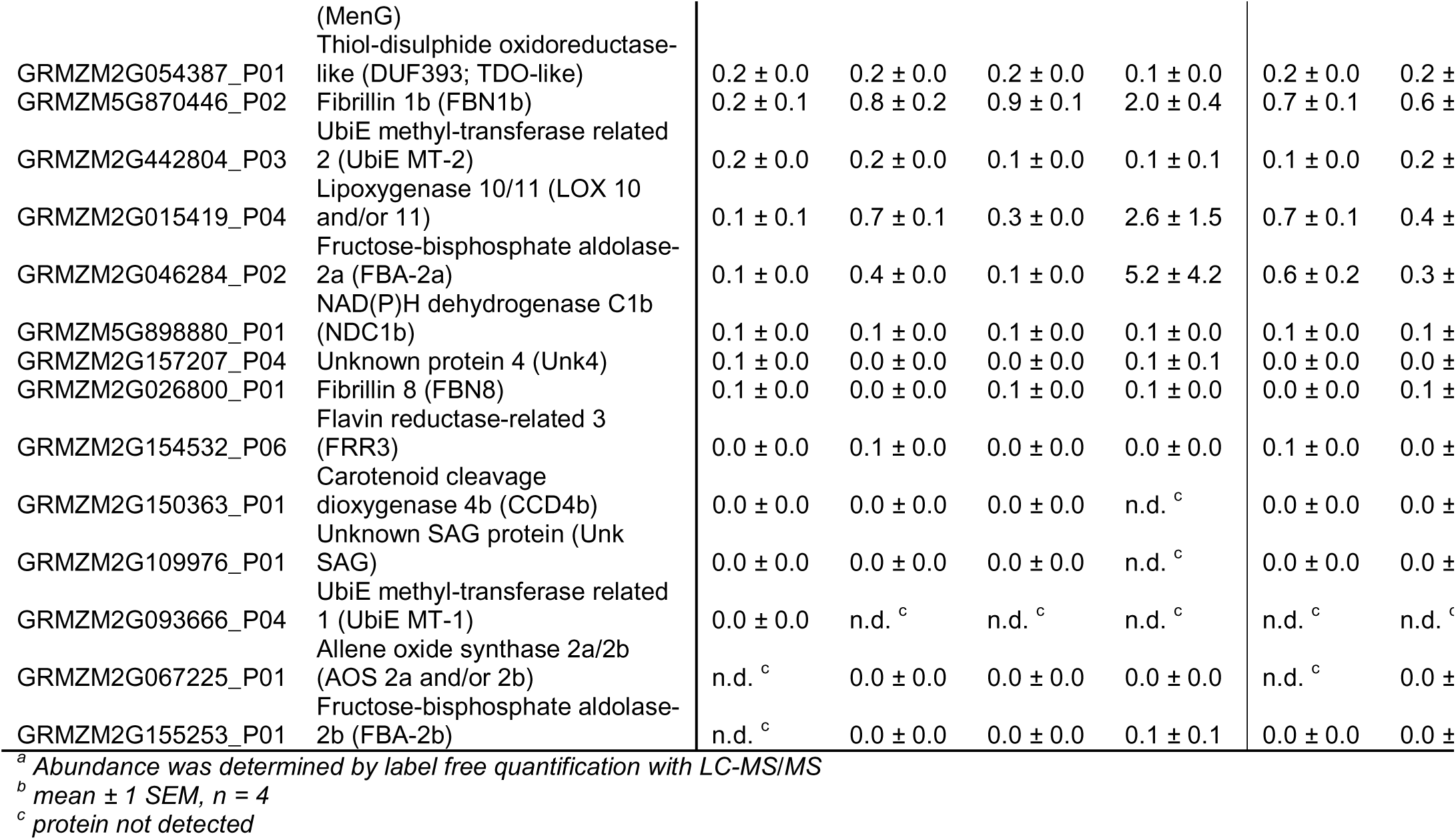
The quantitative *Z. mays* plastoglobule proteome.

Notably, the relative proportions of key protein groups were comparable to what has been reported in *A. thaliana* plastoglobules. For example, FBNs comprised *ca.* 47 – 61 % of the protein mass of maize plastoglobules compared to 42 – 53 % of *A. thaliana* plastoglobules, while ABC1Ks comprised 5-19% in *A. thaliana* plastoglobules and 9 – 12 % in *Z. mays* plastoglobules (**Figure 4B, Supp Table S8**) (Lundquist et al., 2012; Espinoza-Corral et al., 2021). However, the proportions of some proteins were very distinct from that of *A. thaliana* plastoglobules. It was especially striking that a single protein, ZmFBN4, accounts for over 50% of the plastoglobule proteome (**Table 2**). The *A. thaliana* homolog, AtFBN4, is also the most abundant protein of *A. thaliana* plastoglobules, but accounts for a mere 10% of the proteome mass (Espinoza-Corral et al., 2021). Also conspicuous was the reduced proportion of the Plastoglobular Protein of 18 kD (ZmPG18), Fructose bis-phosphate Aldolases and UbiE methyl transferase-like protein which collectively amounts to only 0.7% of protein mass in *Z. mays*, but over 16% in *A. thaliana* (**Table 2**). Thus, while the quantitative proteomes of *Z. mays* and *A. thaliana* are broadly very similar, abundance of some specific proteins are markedly different, potentially related to lineage-specific roles for the plastoglobules.

Accumulation patterns of each of the 38 plastoglobule proteins were plotted by heatmap, revealing changes in abundance over the time course in plastoglobule and thylakoid samples (**Figure 4C**). Nine proteins showed peak plastoglobule abundance at 17 dWW and in most cases quickly declined upon re-watering. These proteins included ZmFBN1a, 1b and 2, Fructose-bisphosphate aldolase-2a and -2b, and Lipoxygenase 10/11 (**Figure 4C**). The elevated accumulation of the ZmFBNs and ZmLOX10/11 at 17 dWW is consistent with their hypothesized roles in regulation of plastoglobule abundance and stress tolerance, particularly in support of JA biosynthesis (Youssef et al., 2010; Lundquist et al., 2012; Borrego and Kolomiets, 2016; Lundquist and Espinoza-Corral, 2022). In contrast, five other proteins showed peak plastoglobule abundance at 7 dWW, including ZmFBN4, ZmPGM48, ZmPES1, and ZmUbiE- MT1. ZmUbiE-MT2 was also abundant at 7 dWW, but peaked at a slightly higher nLFQ intensity at the subsequent time point, 12 dWW. These proteins also tended to show lowest abundance at the 17 dWW time point. It is notable that ZmPGM48 and ZmPES1 decline during stress because these two proteins are associated with promotion of leaf senescence (Lippold et al., 2012; Bhuiyan et al., 2016; Bhuiyan and van Wijk, 2017) and were induced at the plastoglobule in high-light stressed *A. thaliana* (Espinoza-Corral et al., 2019). Furthermore, the Senescence-associated gene of Unknown Function (ZmUnkSAG) was not detected with isolated plastoglobules at 17 dWW, further indicating that the maize plants are not initiating a senescence process under our stress treatment regime. We note that ZmUnkSAG was found on plastoglobules at the earliest time point, albeit at very low levels, while the stress-induced plastoglobule protein ZmELT4, was not detected at any time point.

The most striking effects on the plastoglobule proteome were seen with ZmFBN1a and ZmFBN1b (**Figure 4C**, **Table 2**). These proteins were statistically significantly increased at the plastoglobule during stress, and peaked at 17 dWW, before partially recovering during the re- watering period. Notably, the Protein of Unknown Function 4 (ZmUnk4) closely tracked the accumulation of ZmFBN1a and ZmFBN1b although the changes were not statistically significant, possibly due to its lower accumulation levels. Also striking, was the appearance of two Fructose bis-phosphate aldolase isoforms (FBAs) at the plastoglobule under stress, particularly at 17 dWW when it accounted for over 5% of the plastoglobule proteome (**Figure 4C**, **Table 2**). While multiple FBA isoforms are also associated with *A. thaliana* plastoglobules (Vidi et al., 2006; Ytterberg et al., 2006; Espinoza-Corral et al., 2021), their accumulation is not dependent on the presence of stress, showing comparable levels under both unstressed and high light-stressed conditions (Espinoza-Corral et al., 2021).

Evidence for partial re-mobilization could be seen for several of the FBN proteins, namely FBN4 and the FBN7 paralogs, which declined steadily at the plastoglobule over the course of the stress treatment along with a corresponding increase in levels at the thylakoids over the same time period (**Figure 4D**, **Table 2**). Similarly, the accumulation patterns of the Fructose Bisphosphate Aldolase 2 paralogs, FBA2a and FBA2b, suggested their remobilization from thylakoids to plastoglobules specifically at the peak stress time point, 17 dWW (**Table 2, Supp Fig S5**). Their accumulation in plastoglobule samples spiked at this time point (although the biological variability at the plastoglobules was substantial), which coincided with a clear drop in accumulation at the thylakoid at the same time point. Importantly, the accumulation patterns of the FBN and FBA proteins was reversed within one day of re-watering. Although we cannot determine how these proteins were behaving in the stroma, since stromal samples were not collected or analyzed, the clear patterns of both FBA2 proteins suggest that a *bona fide* pool of FBA2a and FBA2b exists on thylakoid and plastoglobule, and that they are re-directed to the plastoglobules under the stress in a reversible fashion.

### Plastoglobules are comprised of galacto- and sulfo-lipids with acyl chain compositions unique to that of the thylakoids

The membrane surface of plastoglobules has been shown to be contiguous with the outer leaflet of the thylakoid membrane bilayer and thus is presumably comprised of a similar galacto- and sulfo-lipid composition. However, this has not been conclusively demonstrated in the literature.

We sought to profile the polar lipid composition of plastoglobules and their dynamic changes under our water-deficit stress. To do so, we employed thin layer chromatography for lipid separation followed by GC-FID analysis of saponified acyl groups for accurate quantification and acyl chain identification (Wang and Benning, 2011). Consistent with the literature (Block et al., 1983; Douce and Joyard, 1996; Dormann and Benning, 2002), thylakoids were comprised primarily of monogalactosyl diacylglycerol species (MGDG) and digalactosyl diacylglycerol species (DGDG) in a roughly 2:1 ratio, with smaller amounts of sulfoquinovosyl diacylglycerol (SQDG; *ca.* 4 mol%) and phosphatidylglycerol (PG; *ca.* 8 mol%) also present (**Figure 5A, Supp Table S9)**. These ratios changed only slightly over the time course, with SQDG declining significantly during the water-deficit and DGDG commensurately increasing under water-deficit; both trends were reversed between 1 and 3 days after re-watering. While plastoglobules were also rich in galactolipids, they skewed towards a somewhat greater quantity of MGDG and lacked any PG. Because of the larger variation among replicates of the plastoglobules, it was difficult to discern changes in polar lipid levels over the time course.

**Figure 5.**
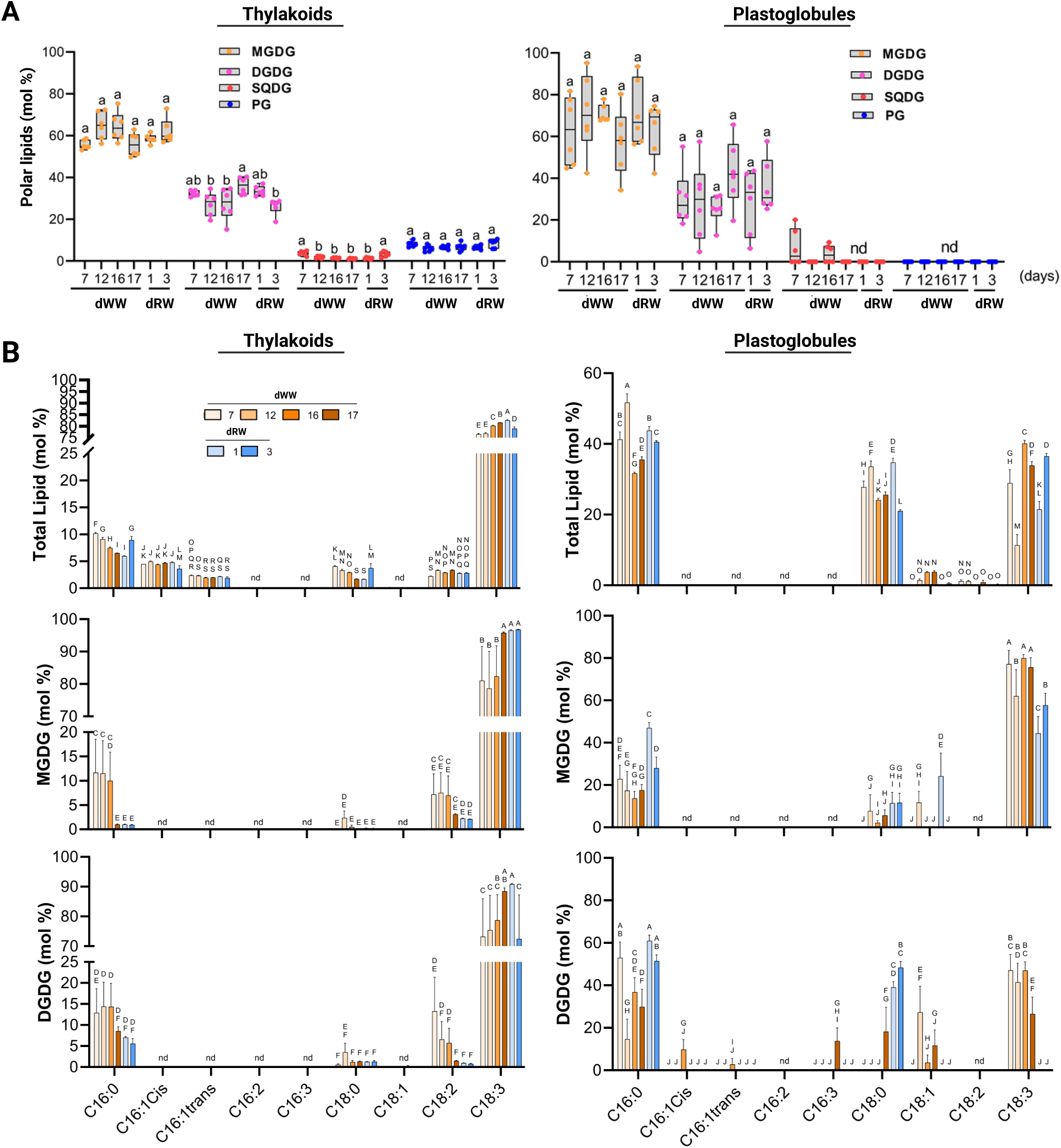
Lipidome profiling of isolated *Z. mays* plastoglobules and thylakoids during the water- deprivation-and-recovery time course. **A)** Polar lipids were separated by thin layer chromatography, scraped from the plates, saponified and analyzed by gas-liquid chromatography with flame ionization detection. Relative molar ratios of each polar lipid class are plotted as box-and-whisker plots with individual data points overlaid. Statistically significant groupings between time points (and within a polar lipid class) are indicated by letter codes using two-way ANOVA, p < 0.001, n = 6 individual plants. Monogalactosyl diacylglycerol (MGDG), digalactosyl diacylglycerol (DGDG), sulfoquinovosyl diacylglycerol (SQDG), and phosphatidylglycerol (PG). **B)** Relative molar ratios of the acyl chains from total lipid extracts and scraped spots of MGDG and DGDG of isolated thylakoids (left) and isolated plastoglobules (right) at each time point. Statistically significant groupings between time points (and within a polar lipid class) are indicated by letter codes using two-way ANOVA, p < 0.001, n = 6 individual plants.

We observed highly distinct acyl chain compositions amongst the lipids of plastoglobule and thylakoids. Whereas total lipid extracts of thylakoid were predominantly comprised of C18:3 fatty acid, comprising *ca.* 85 mol%, consistent with a preponderance of C18:3 fatty acid reported in the literature (Hiller and Goodchild, 1981; Browse et al., 1986), those of the plastoglobules were comprised of roughly equal proportions of C18:3 and C16:0, highlighting a striking enrichment of saturated fatty acyl chains (**Figure 5B, Supp Table S10**). When looking at the acyl chain composition of individual polar lipids this pattern was maintained. MGDG and DGDG of the thylakoid were comprised of a minimum of 80 mol% C18:3 at all time-points, while SQDG and PG were comprised of significant amounts of C16:0 in addition to C18:3 (**Figure 5B, Supp Table S10**). This is in sharp contrast to the plastoglobules, in which saturated acyl chains comprise a substantial proportion of the MGDG and DGDG lipids at all time points. Moreover, the SQDG of the plastoglobules was found to only contain C16:0 or C18:0 saturated acyl chains (**Supp Table S10**). Also striking is the observation that whereas thylakoids progressively reduced their levels of saturated acyl chains during the progression of the stress, the plastoglobules showed the reverse pattern, with an increase in their proportion of saturated acyl chains. For example, DGDG in the plastoglobules progressed from *ca*. 50 mol% saturated acyl chains at the 7 dWW to 100 mol% saturated acyl chains by the final time point. It remains to be determined whether this reflects a transfer of galactolipid (with saturated acyl chains) from thylakoid to plastoglobule under the water-deprivation stress. Collectively, we conclude that plastoglobules contain a *bona fide* population of galactolipids with a highly distinct acyl chain composition enriched in saturated fatty acid.

### Prenyl-lipid profiling of thylakoids and plastoglobules

Plastoglobules and thylakoids are rich stores of various prenyl-lipid compounds, such as carotenoids and quinones, involved in photosynthesis and/or reactive oxygen scavenging (Lundquist et al., 2013; van Wijk and Kessler, 2017). We profiled the prenyl-lipid composition of our isolations using ultra-reversed phase (C30 column) high performance lipid chromatography (HPLC) with photodiode array detection (PDA) (Fraser et al., 2000). To provide an appropriate normalization across the samples we quantified thylakoid prenyl-lipid species relative to total protein, and plastoglobule prenyl-lipid species to a 200 µL volume of plastoglobule suspension at 1.0 OD_700_. The thylakoids and plastoglobules were rich in cumulative levels of plastoquinone- 9 and plastoquinol-9 (PQ-9 and PQH2, respectively; oxidized and reduced forms) under both water-deficit and recovery conditions with approximately 18% and 94% of the prenyl-lipid mass of thylakoids and plastoglobules, respectively (**Figure 6**). Because we did not take precautions to prevent artefactual oxidation of PQH2-9 to PQ-9 we do not place any significance on the ratio between these two compounds and instead consider them as a single form [PQ(H_2_)-9]. There is a substantially higher proportion of PQ(H_2_)-9 in *Z. mays* plastoglobules than has been reported in *A. thaliana* (*ca.* 60 mol % (Lundquist et al., 2013; Espinoza-Corral et al., 2021). The relative abundance of prenyl-lipids detected in both thylakoids and plastoglobules were unchanged during the time course. The lack of apparent remobilization of carotenoid pigments from thylakoids to plastoglobules - as has been observed under light stressed *A. thaliana* (Espinoza-Corral et al., 2021) – indicate that dismantling of the thylakoid has not been initiated under our conditions, consistent with our proteomics results above.

**Figure 6.**
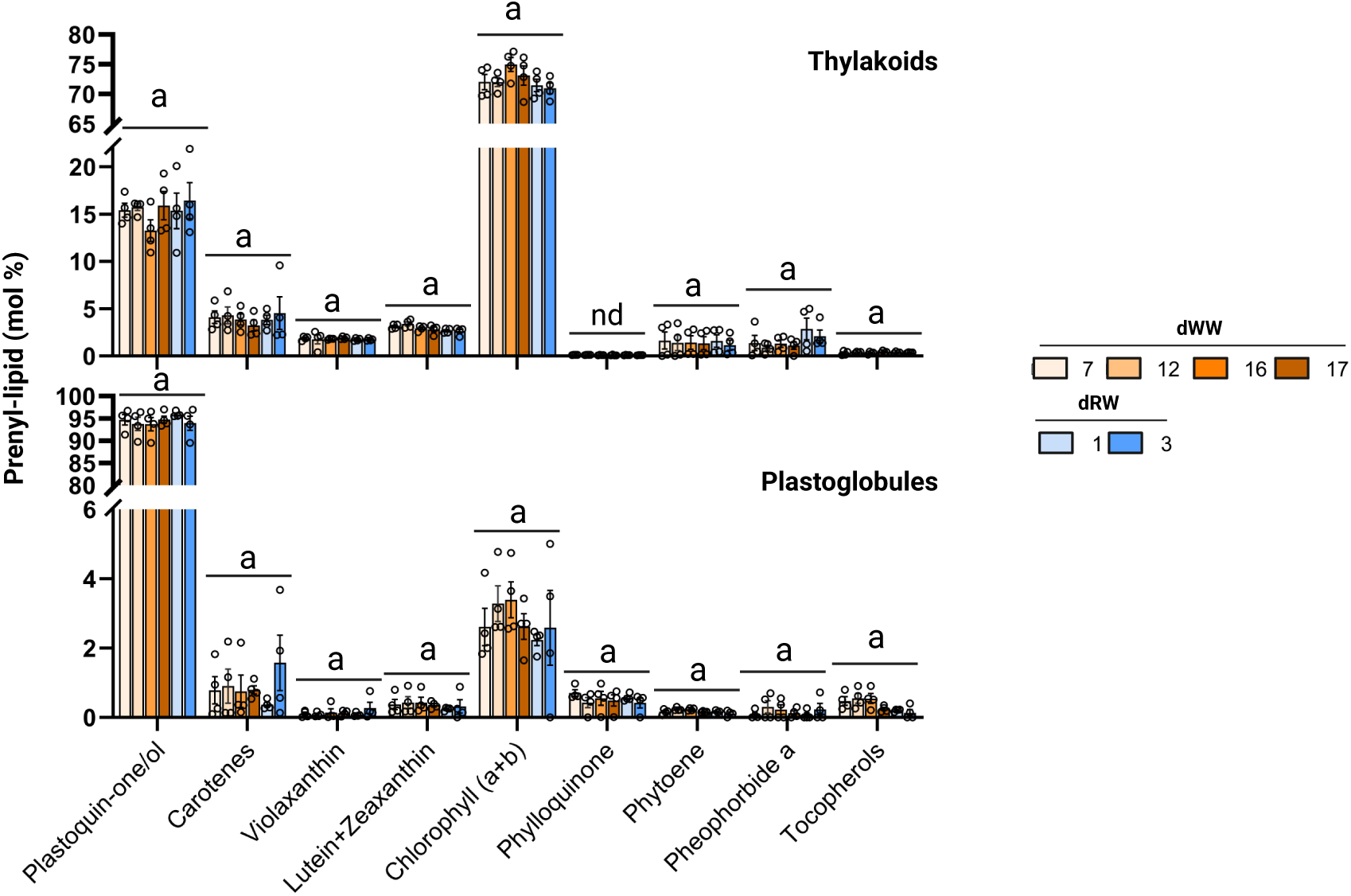
Prenyl-lipid profiles of isolated Z. mays plastoglobules and thylakoids during the water-deficit-and-recovery time course. Relative molar ratios of each prenyl-lipid class are plotted as bar plots, mean ± 1 s.e.m., with individual data points overlaid as open circles, n = 4 biological replicates. Reduced and oxidized forms of plastoquinone are quantified together, as well as chlorophylls a and b, and lutein and zeaxanthin.

### LC-MS/MS lipidomic analysis of plastoglobules and thylakoids

Liquid chromatography connected in-line with tandem mass spectrometry (LC-MS/MS) can provide robust identification of lipid species through accurate mass measurement of intact ions and analysis of their fragmentation patterns. We employed LC-MS/MS to attempt to identify additional lipid species in our thylakoid and plastoglobule samples. In the absence of standards for quantification, we were not able to quantify relative proportions of different compounds. However, ion intensities could be used to estimate relative changes of each lipid over the time course. A total of 7,028 distinct molecular species (*i.e.* unique m/z and retention time values) were found across the thylakoid samples, and 6,804 across the plastoglobule samples (**Supp Table S11 and S12**). Of these, 109 (thylakoids) and 90 (plastoglobules) could be annotated as specific lipid species by matching accurate mass measurements and/or fragmentation patterns, accounting for *ca.* 15% and 10%, respectively, of the total MS1 ion intensity of each sample. Most of the annotated lipids were found in both sub-compartments.

We first note that neutral lipids in mono-, di- and tri-acylglycerol forms were found in both sub-compartments (**Figure 7, Supp Table S11 and S12**). Notably, monoacylglycerols (MAGs) and diacylglycerols (DAGs) were found in about 10-fold lower proportions in plastoglobules than in thylakoids. This striking difference suggests that detectable levels in the plastoglobules may derive from residual thylakoid contamination and that they are, in fact, excluded from plastoglobules. This contrasts with the triacylglycerols (TAG) which accounted for a much larger proportion of the total ion intensity of plastoglobules than that of thylakoids. Thirty-two distinct TAG species were annotated in the plastoglobule and thirty-three in thylakoids, including most combinations of 16 and 18 C acyl groups (**Figure 7B**, **Table 3**). We additionally found low levels of TAG with 56:0, 56:1 and 56:2 (where the number preceding the colon indicates the collective number of acyl carbons and the number following the colon indicates the collective number of double bonds), indicating that some arachidic acid (a saturated 20 C acyl group) is incorporated into the plastoglobule TAG.

**Figure 7.**
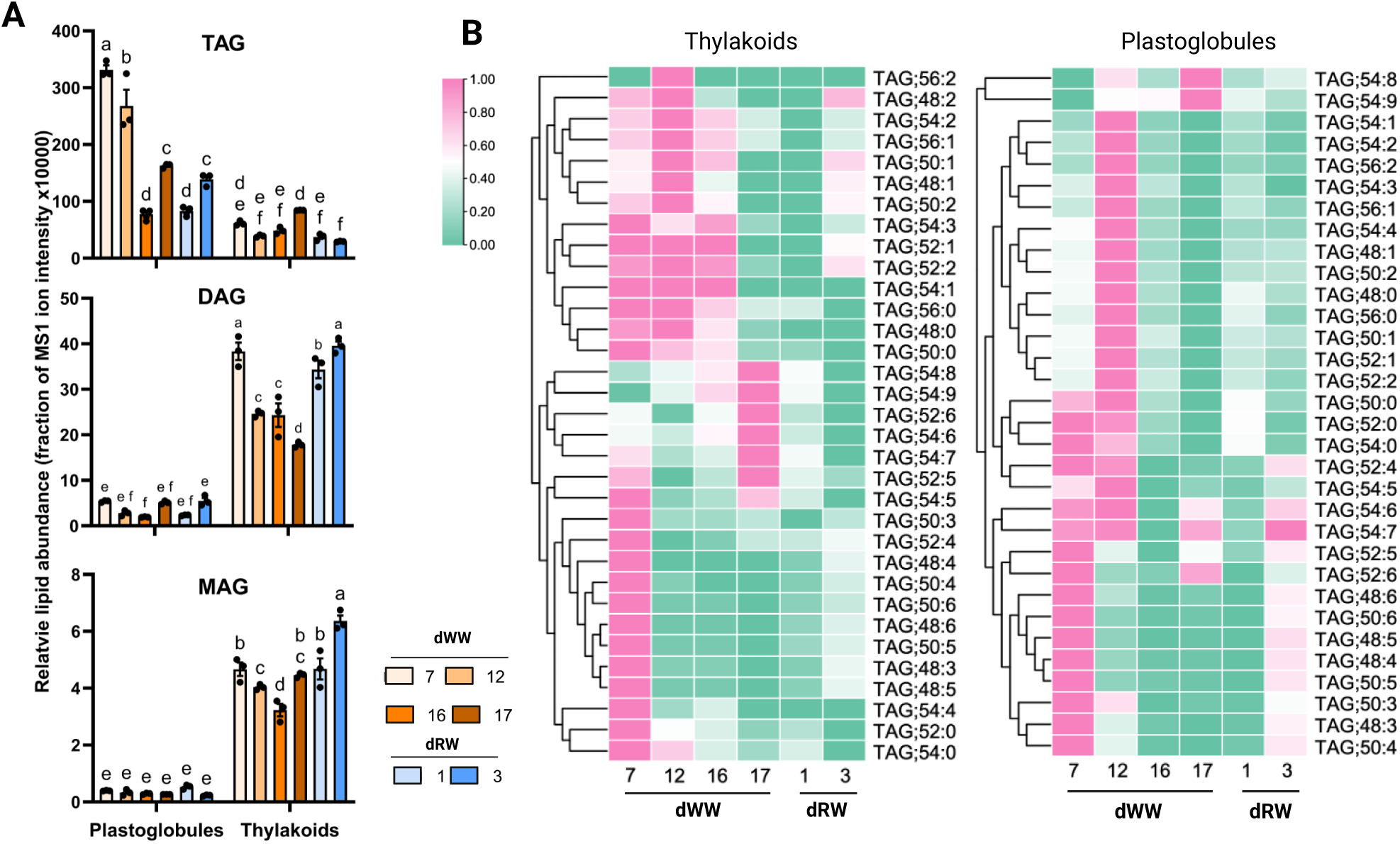
Relative levels of neutral lipids in *Z. mays* plastoglobules and thylakoids. Bar charts presenting the mean cumulative MS1 ion intensities of each class of neutral lipid. Mean ± 1 s.e.m., n = 3 biological replicates. with black dots representing individual data points. TAG, triacylglycerol ; DAG, diacylglycerol; MAG, monoacylglycerol.

**Table 3.** Acyl composition of triacylglycerol species (TAGs) in plastoglobules at each time point.

| Acyl composition | 7 dWW | 12 dWW | 16 dWW | 17 dWW | 1 dRW | 3 dRW |
| --- | --- | --- | --- | --- | --- | --- |
| 48:0<sup>a</sup> | 1.6 ± 0.1<sup>b</sup> | 4.9 ± 1.2 | 4.6 ± 0.2 | 1.4 ± 0.1 | 6.0 ± 0.7 | 2.5 ± 0.5 |
| 48:1 | 0.8 ± 0.2 | 3.9 ± 0.2 | 1.5 ± 0.5 | 0.3 ± 0.0 | 1.7 ± 0.6 | 1.2 ± 0.3 |
| 48:3 | 5.3 ± 0.2 | 1.3 ± 0.0 | 1.3 ± 0.3 | 0.5 ± 0.0 | 1.1 ± 0.1 | 4.0 ± 0.1 |
| 48:4 | 4.7 ± 0.2 | 0.4 ± 0.0 | 0.7 ± 0.1 | 0.3 ± 0.0 | 0.5 ± 0.0 | 3.8 ± 0.2 |
| 48:5 | 8.9 ± 0.4 | 0.5 ± 0.1 | 0.8 ± 0.1 | 0.5 ± 0.0 | 0.7 ± 0.0 | 6.6 ± 0.8 |
| 48:6 | 10.3 ± 0.4 | 1.3 ± 0.3 | 1.7 ± 0.1 | 1.1 ± 0.0 | 1.5 ± 0.2 | 6.7 ± 0.2 |
| 50:0 | 1.6 ± 0.1 | 2.8 ± 0.8 | 2.7 ± 0.1 | 0.8 ± 0.1 | 3.9 ± 0.4 | 1.3 ± 0.3 |
| 50:1 | 1.6 ± 0.5 | 6.2 ± 0.7 | 5.2 ± 1.1 | 1.0 ± 0.1 | 4.7 ± 0.6 | 2.3 ± 0.3 |
| 50:2 | 1.4 ± 0.3 | 5.6 ± 0.5 | 3.2 ± 0.7 | 0.8 ± 0.1 | 3.3 ± 0.8 | 2.0 ± 0.4 |
| 50:3 | 3.9 ± 0.1 | 2.4 ± 0.1 | 1.9 ± 0.1 | 1.0 ± 0.0 | 1.9 ± 0.3 | 3.4 ± 0.1 |
| 50:4 | 4.5 ± 0.2 | 1.3 ± 0.2 | 1.2 ± 0.2 | 0.5 ± 0.0 | 1.2 ± 0.0 | 4.0 ± 0.1 |
| 50:5 | 6.0 ± 0.3 | 0.3 ± 0.1 | 0.6 ± 0.1 | 0.2 ± 0.0 | 0.4 ± 0.1 | 4.2 ± 0.3 |
| 50:6 | 9.2 ± 0.4 | 0.5 ± 0.1 | 0.9 ± 0.1 | 0.5 ± 0.0 | 0.7 ± 0.1 | 5.1 ± 0.1 |
| 52:0 | 2.8 ± 0.1 | 3.2 ± 1.2 | 2.7 ± 0.4 | 0.9 ± 0.1 | 4.8 ± 0.3 | 1.1 ± 0.1 |
| 52:1 | 0.6 ± 0.2 | 2.2 ± 0.3 | 1.7 ± 0.2 | 0.4 ± 0.0 | 1.8 ± 0.3 | 0.8 ± 0.1 |
| 52:2 | 1.4 ± 0.5 | 6.6 ± 0.7 | 4.7 ± 1.0 | 0.9 ± 0.2 | 4.4 ± 0.05 | 1.5 ± 0.3 |
| 52:4 | 4.1 ± 0.1 | 4.4 ± 0.5 | 2.7 ± 0.1 | 1.5 ± 0.0 | 3.2 ± 0.1 | 5.1 ± 0.3 |
| 52:5 | 5.3 ± 0.2 | 2.2 ± 0.2 | 3.1 ± 0.3 | 4.0 ± 0.1 | 3.9 ± 0.5 | 5.8 ± 0.4 |
| 52:6 | 8.4 ± 0.4 | 2.3 ± 0.3 | 6.7 ± 0.4 | 13.0 ± 0.2 | 5.2 ± 0.7 | 6.6 ± 0.2 |
| 54:0 | 2.8 ± 0.1 | 2.4 ± 0.9 | 2.5 ± 0.4 | 0.8 ± 0.1 | 4.5 ± 0.2 | 1.2 ± 0.1 |
| 54:1 | 0.2 ± 0.0 | 1.2 ± 0.3 | 0.6 ± 0.1 | 0.1 ± 0.0 | 0.7 ± 0.2 | 0.2 ± 0.0 |
| 54:2 | 0.3 ± 0.1 | 2.0 ± 0.2 | 1.2 ± 0.2 | 0.3 ± 0.0 | 1.2 ± 0.2 | 0.4 ± 0.0 |
| 54:3 | 1.2 ± 0.6 | 6.8 ± 2.6 | 3.8 ± 1.0 | 0.7 ± 0.2 | 3.4 ± 0.6 | 0.8 ± 0.1 |
| 54:4 | 1.7 ± 0.3 | 6.9 ± 1.9 | 3.2 ± 0.7 | 0.8 ± 0.1 | 2.9 ± 0.4 | 1.6 ± 0.1 |
| 54:5 | 2.1 ± 0.1 | 5.3 ± 1.2 | 1.9 ± 0.2 | 1.4 ± 0.0 | 2.1 ± 0.1 | 2.4 ± 0.0 |
| 54:6 | 3.7 ± 0.1 | 5.2 ± 0.2 | 3.3 ± 0.2 | 4.5 ± 0.1 | 3.9 ± 0.5 | 6.2 ± 0.2 |
| 54:7 | 2.7 ± 0.1 | 3.5 ± 0.3 | 4.6 ± 0.2 | 5.1 ± 0.1 | 5.0 ± 0.9 | 7.1 ± 0.3 |
| 54:8 | 1.4 ± 0.1 | 5.1 ± 0.8 | 8.9 ± 0.3 | 15.9 ± 0.4 | 8.5 ± 1.2 | 6.5 ± 0.2 |
| 54:9 | 0.7 ± 0.0 | 5.3 ± 1.2 | 19.5 ± 0.8 | 40.3 ± 0.5 | 13.3 ± 2.1 | 4.3 ± 0.3 |
| 56:0 | 0.3 ± 0.0 | 1.1 ± 0.4 | 0.9 ± 0.1 | 0.3 ± 0.0 | 1.3 ± 0.2 | 0.4 ± 0.1 |
| 56:1 | 0.4 ± 0.1 | 1.8 ± 0.2 | 1.2 ± 0.1 | 0.3 ± 0.0 | 1.5 ± 0.4 | 0.5 ± 0.1 |
| 56:2 | 0.2 ± 0.1 | 0.9 ± 0.1 | 0.6 ± 0.0 | 0.3 ± 0.0 | 0.6 ± 0.1 | 0.3 ± 0.0 |

<sup>a</sup> The cumulative composition of all three acyl groups are described, with the total number of C atoms represented in the number preceding the colon and the total number of double bonds following the colon.
<sup>b</sup> Percent fraction of total TAG MS1 ion intensity given as mean ± 1 s.e.m, n = 3 biological replicates.

Surprisingly, the proportion of TAG relative to total lipid in the plastoglobules (measured as MS1 ion intensity of each sample) declined under drought stress, contrasting with the striking increase of plastoglobule-associated TAG during heat stress documented in our companion in paper (Devadasu, et al. unpublished). This indicates that the increase in plastoglobule size and abundance is not primarily driven by TAG accumulation under water-deficit stress **(Supp Tables S11 and S12**).

Prior to the stress treatment, the most abundant TAG species in plastoglobules were 48:6, 48:5, and 50:6 (**Table 3**). However, under stress the relative levels of these species decline substantially and are replaced with longer chain acyl groups, most notably 54:9 and 54:8. The quantitative change in levels of these lipids is quite dramatic, with the relative amount of the longer chain TAG species increasing from about 1% each of the total TAG amount, to over 40% in the case of 54:9, and over 15% in the case of 54:8. Importantly, while TAG is also identified in our isolated thylakoid samples, the relative proportions of the species largely mirror that of the plastoglobule samples at each time point (**Figure 7B**, **Supp Table S11**), indicating that the TAG identified in thylakoids is derived primarily from plastoglobule material that co-isolates with the thylakoids. This is reasonable since the bilayer membranes of the thylakoid could not effectively store large quantities of TAG.

The highest MS1 ion intensity in plastoglobules was from PQ-9, consistent with the high levels found with our HPLC-PDA analysis (**Figure 6**). In addition, we also annotated peaks as plastochromenol (PC-8) and possibly plastochromanol (PCH_2_-8) in our samples, which could not be confidently identified by HPLC-PDA due to the lack of authentic standards. These are the oxidized and reduced forms, respectively, of the chromanol form of PQH_2_-9, synthesized from PQH_2_-9 by plastoglobule-localized Tocopherol Cyclase, *i.e.* VTE1 (Szymańska and Kruk, 2008; Szymanska and Kruk, 2010; Kruk et al., 2014). Notably, the MS1 ion intensities of PC-8, and the possible PCH_2_-8, were several-orders of magnitude lower than PQ-9. Although the relative ionization efficiencies of these species are unclear, this indicates the ratio of PC-8 to PQ-9 may be skewed towards PQ-9 in *Z. mays* plastoglobules which would contrast sharply with *A. thaliana* plastoglobules in which they were found in roughly equal proportions (Lundquist et al., 2013).

Notably, with LC-MS/MS we were also able to identify low levels of phosphatidylglycerol and SQDG in our plastoglobule samples, contrasting with the GC-FID analysis where they were typically below the detection limit. However, the acyl composition of both PG and SQDG were similar to that in the thylakoids (**Supp Tables S11 and S12**). This observation, combined with their very low levels in the plastoglobules, suggests they may represent contamination from the thylakoids. In contrast, the LC-MS/MS analysis supports the apparent enrichment of saturated acyl groups on MGDG and DGDG in plastoglobules that we had observed by GC-FID (**Figure 5**). Thus, we conclude that the *Z. mays* plastoglobule surface is comprised of MGDG and DGDG, but not SQDG or PG. We also considered whether lyso- forms of MGDG or DGDG may be present in *Z. mays* plastoglobules, which may be more favorable on the tightly curved plastoglobule surface, however we could not identify evidence of such lipids by LC-MS/MS of thylakoids or plastoglobules.

### LC-MS/MS evidence for multiple plastoquinone derivatives in the maize plastoglobules

Various derivatives of PQ(H_2_)-9 have been described in the literature such as PQ(H_2_)-9 that is hydroxylated at one of the double bonds on its solanesyl side chain, or that is hydroxylated and then subsequently acylated at this hydroxyl group (Kruk et al., 1998; Kruk and Strzałka, 1998). To distinguish these derivatives from the canonical (*i.e.*, unmodified) PQ(H_2_)-9, a nomenclature has been established assigning a different letter for each PQ derivative. In this way, the unaltered PQ(H_2_)-9 is referred to as PQ(H_2_)-A, those hydroxylated on the prenyl tail as PQ(H_2_)- C, and those hydroxylated and subsequently acylated on the prenyl tail as PQ(H_2_)-B (Griffiths et al., 1966; Barr et al., 1967; Susanto et al., 2025). Note that a class of PQ derivatives originally referred to as PQ(H_2_)-D were later determined to be alternative isomers of PQ(H_2_)-C (*i.e.*, hydroxylated at different position on the prenyl tail), and the name PQ(H_2_)-D was subsequently retired (Barr et al., 1967; Das et al., 1967). Despite the discovery of these derivatives over 50 years ago, their site of accumulation and functional significance remains unclear.

In addition to the above PQ(H_2_)-9 derivatives, a novel PQ(H_2_)-9 derivative, in which an acyl group is attached to one of the hydroxyl groups of the PQH_2_ head group, has recently been identified in cyanobacteria (Ishikawa et al., 2023; Kondo et al., 2023; Mori-Moriyama et al., 2023). We have named this novel PQ(H_2_)-9 derivative as PQH_2_-E and shown that it, and the other PQ(H_2_)-9 derivatives, accumulate in large quantities in the cyanobacterial lipid droplets known as cyanoglobules (Susanto et al., 2025).

With our LC-MS/MS investigation in *Z. mays* we have annotated multiple members of each derivative class in both the plastoglobule and thylakoid samples (**Figure 8, Supp Table S11 and S12**). The MS1 ion intensities of PQ(H_2_)-9 derivatives were substantial and were among the largest intensities across all annotated lipids. This suggests that these compounds are prominent components of *Z. mays* plastoglobules and are well represented by PQ-B, PQH2-E, and PQ(H_2_)-C forms (**Figure 8A**).

**Figure 8.**
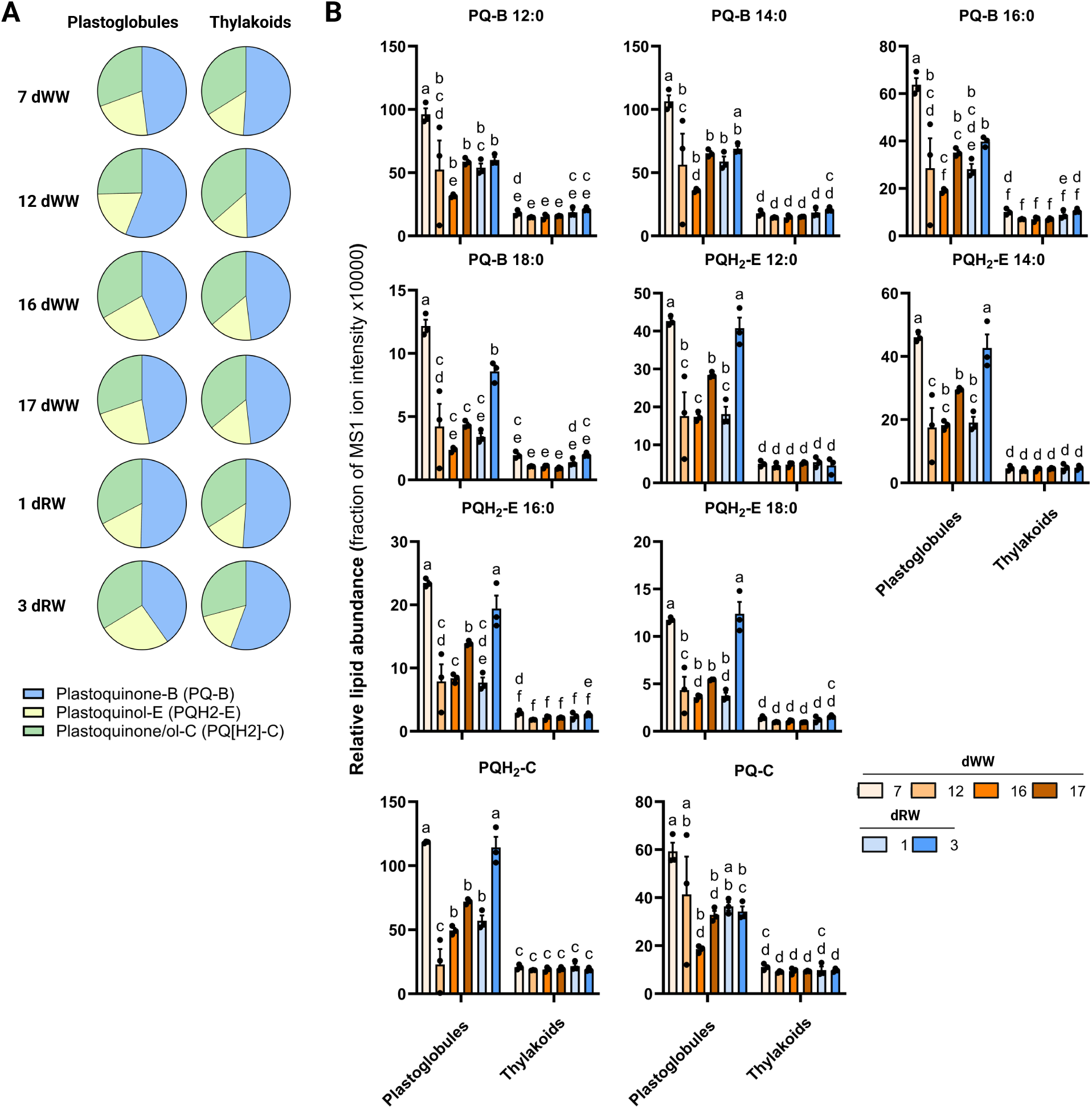
Relative levels of plastoquinone derivatives in Z. mays plastoglobules and thylakoids. **A,** Pie chart of mean cumulative MS1 ion intensities of each class of PQ derivatives in isolated plastoglobules and isolated thylakoids at each time point, n = 3 biological replicates. **B,** Bar graphs depicting relative levels of each individual PQ derivative identified in plastoglobules and thylakoids. Mean ± 1 s.e.m with black dots representing individual data points, n = 3 biological replicates.

Strikingly, PQ-B species were only annotated with the oxidized head group, never the reduced PQH_2_-B form. In addition, only saturated acyl groups were identified within the PQ-B and PQH_2_- E series, comprising the 12:0, 14:0, 16:0 and 18:0 forms. The relative ion intensities indicated that the 12:0 form was the dominant acyl group on both PQ-B and PQH_2_-E. We also note that only a single isomer each of PQ-C and PQH_2_-C were found (**Supp Figure S6**). Hydroxylation could in principle occur at any one of the double bonds of the prenyl tail, which could be expected to separate on our LC-MS system. That we find only one such isomer of each of the oxidized and reduced forms suggests that hydroxylation of the prenyl tail, generating PQ(H_2_)-C, may be enzymatically catalyzed. As we demonstrate above with our proteomic analyses, several candidate oxidoreductases are found associated with the plastoglobules. Relative proportions of each PQ derivative class were somewhat depleted in the plastoglobules under the drought stress treatment, possibly due to a dilution effect as other lipids increase in accumulation, although levels were generally stable across the time course in thylakoids (**Figure 8B, Supp Tables S11 and S12**).

In our previous investigation of cyanoglobules of *Synechocystis* sp. PCC 6803 we also annotated a novel phylloquinone with an acylation at the headgroup. However, we could not detect any acylated phylloquinone in plastoglobules or thylakoids of *Z. mays*. In sum, we onclude that the PQ derivatives, PQ-B, PQ(H_2_)-C, and PQH_2_-E, are prevalent in both plastoglobules and thylakoids of *Z. mays* and their levels remain generally stable over the course of the water-deficit stress treatment.

## DISCUSSION

In the present work we have described the protein and lipid composition of *Z. mays* plastoglobules and delved into the effects of a water-deficit stress and recovery treatment on their composition. Drought can profoundly impact photosynthesis in plants, leading to alterations in membrane fluidity and thylakoid membrane disorganization, and has frequently been associated with proliferation of plastoglobules (Munne-Bosch and Alegre, 2004; Grigorova et al., 2012; Shao et al., 2016; Carrera et al., 2021; Chen et al., 2022; Aliyeva et al., 2023). It is reasonable to speculate that the increase in both the size and number of plastoglobules is linked to the accumulation of products resulting from thylakoid membrane turnover. We demonstrate that water-deficit coincided with an increase in the size and abundance of plastoglobules (**Figure 3**). Although we could not discern clear effects on thylakoid ultrastructure in this investigation, evidence of remodeling was apparent from proteomic and lipidomic analysis, including several of the FBNs. As expected, the proliferation of plastoglobules corresponded to a discernable increase in several plastoglobule proteins, such as FBN1a, FBN1b, ABC1K9, and others. This determination is dependent on our investigation of the total leaf proteome, and we can reasonably anticipate that other, lower abundant, members of the plastoglobule proteome also increased, although undetectable at the whole leaf level. Similarly, the proliferation of plastoglobules likely corresponds to an increase in the levels of many of the associated lipids. However, in the absence of lipidomic investigation at the total leaf level, this remains uncertain.

We demonstrate that TAG and various PQ derivatives represent major components of the *Z. mays* plastoglobules. While these lipids remain prevalent under the stress, we were surprised to see that their abundance relative to total plastoglobule lipid (*i.e.*, total MS1 ion intensity) is reduced *ca.* 2-3-fold in all cases, indicating that biosynthesis of these lipids, *per se*, is not a primary driver of plastoglobule proliferation under our water-deficit stress. Our results do not identify obvious candidate lipids that may be accumulating in plastoglobules under the water-deficit stress and driving their proliferation. One possibility is that proliferation of plastoglobules entails increased accumulation of all plastoglobule lipids, which would manifest with at least roughly equal proportions of the lipid composition under both stressed and unstressed conditions. Alternatively, other lipids that are less amenable to our identification methods may

be increasing substantially under stress, such as the fatty acid phytyl esters, which are known to be stored in substantial quantities in *A. thaliana*. Indeed, in our companion paper we have shown that *Z. mays* plastoglobules accumulate the 12:0 FAPE under heat stress (Devadasu, et al. unpublished). In the paragraph below, we note the relatively stable composition of plastoglobule lipid and protein composition in response to stresses and across species. This observation argues for the notion that plastoglobule proliferation is an effect of increased levels of most, or all, plastoglobule lipids, rather than a subset of specific lipids.

The lipidome composition of the plastoglobule has been only incompletely elucidated thus far. Our results demonstrate that the plastoglobule surface is, at least in part, comprised of the galactolipids, MGDG and DGDG. It was surprising to discover that the relative proportions of these two lipid classes are quite comparable to that found in the bulk thylakoid membrane because the relatively tight positive curvature of the plastoglobule surface would be expected to disfavor the presence of the conical MGDG. In fact, we initially considered a possible enrichment of lyso- forms of MGDG or DGDG (*i.e.*, mono-acyl forms of the galactolipids), which would seemingly be better accommodated on the plastoglobule surface. However, our lipidomics failed to detect such lipids. Nonetheless, there was a clear bias towards saturated fatty acids among the plastoglobule MGDG and DGDG when compared to what was seen in the bulk thylakoid.

The compositions of plastoglobules and cyanoglobules have now been identified from *Z. mays*, *Synechocystis* sp. PCC 6803 (Susanto et al., 2025) and *A. thaliana* (Espinoza-Corral et al., 2021), although LC-MS/MS-based lipidomics is still lacking with *A. thaliana* plastoglobules. It has become evident that these compartments harbor broadly similar lipid and protein compositions, largely associated with lipid metabolism and oxidative detoxification. As in *A. thaliana* plastoglobules under a light stress (Espinoza-Corral et al., 2021), the *Z. mays* plastoglobules under the water-deficit stress do not undergo wholesale changes to their lipidome or proteome. Rather, specific, tailored changes are apparent, seemingly related to effects on specific metabolic pathways.

Indeed, we found evidence of recruitment of specific proteins and pathways to the plastoglobules during the stress. Of particular note, we saw a sharp increase in the proportion of LOX 10 and/or 11, AOS 2a and/or 2b, and two isoforms of FBA. The apparent recruitment of these AOS and LOX enzymes holds a striking resemblance to the apparent recruitment of LOX and AOS seen in *A. thaliana* under high-light stress (Espinoza-Corral et al., 2021). In the context of the *A. thaliana*, we proposed that this reflected a senescence-like cell death process that was initiated in response to the stress, a similar explanation may be plausible in the context of the *Z. mays* water-deficit stress. The increase in levels of the FBA isoforms is also notable. Considering our observations under heat stress (Devadasu, et al. unpublished), in which we observed the enrichment of many of the Calvin-Benson cycle enzymes (in addition to FBA), we must emphasize that under the water-deficit stress described here, we only observed the FBA isoforms associated with plastoglobules, not other Calvin-Benson cycle enzymes. The presence of the FBA enzymes is consistent with the routine observation of a subset of the FBA pool associated with *A. thaliana* plastoglobules (Vidi et al., 2006; Ytterberg et al., 2006; Lundquist et al., 2012). Nonetheless, the physiological significance of this localization of FBA remains unclear.

By a wide margin the most highly abundant protein of the plastoglobule proteome was FBN4. Although no functions for FBN4 have been demonstrated, FBN4 has been shown to positively correlate with PQ-9 levels in the plastoglobules of Malus x domestica (Singh et al., 2010). Thus, the high levels of PQ-9 that we observe in the *Z. mays* plastoglobules are notable and lends further credence to a possible role for FBN4 in promoting PQ-9 accumulation in plastoglobules through a yet unknown mechanism. In light of the atypical lipocalin domain of most FBNs, Singh and McNellis have previously proposed that FBN4 may bind and transport PQ-9 (Singh and McNellis, 2011). This is a compelling possibility that is further supported by the observed role for the close homolog, FBN5, in supporting PQ-9 biosynthesis (Kim et al., 2015).

As a C4 grass, the morphological features and functions of M and BS cells of *Z. mays* are divergent. Thus, it is possible that some degree of divergence in composition or function exists between plastoglobules of these two cell types. Indeed, we documented differing morphological responses of plastoglobules in M and BS cells, in which those of M cells uniquely increased in abundance, while plastoglobules of M and BS cells increased in abundance under the stress (**Figure 3**). However, it is important to note that our methodology for isolation of plastoglobules and thylakoids does not allow us to distinguish between these two cell types. Homogenization of the leaf tissue, the first step of our isolation protocol, means any possible heterogeneity among plastoglobules is lost. Thus, we must consider our analyses as a cumulative view across both cell types. The requirement for relatively large quantities of leaf tissue for efficient isolation of plastoglobules makes it difficult to uncover possible heterogeneity, whether between cell types or within the same cell or chloroplast, although such heterogeneity may very well exist.

## MATERIAL & METHODS

### Plant materials, water deficit treatment, and measurement of relative water content

Maize (*Zea mays* ssp. *mays* var. B73) seeds were washed with distilled water twice and kept in the dark overnight. Seeds were then sown in plastic pots (diameterL×Lheight: 9 ×L10 cm) filled with Sure Mix soil and germinated in a growth chamber (Biochambers Reach-in SPC-37, Winnipeg, Canada) with a 16 h photoperiod and day/night temperatures of 27 and 23L °C, respectively. Photosynthetic photon flux density from LED lighting was *ca.* 600LμmolLm^−2^Ls^−1^ and relative humidity was 50–60%. Equal volumes of water (200 mL) were provided to each pot on a weekly basis. Three weeks after germination, when four leaves were fully expanded, a progressive drought stress was imposed by withholding water for a total of 17 days. Imposition of stress and recovery (re-watering) was monitored by measuring the relative water content (RWC) of the fourth or fifth collared leaf on each day of the time course approximately two hours after lights were turned on. Leaf RWC was estimated using the relative turgidity method (Weatherley, 1950). Briefly, the fourth or fifth leaf was harvested and immediately measured for fresh weight (FW). Turgid weight (TW) was then determined after leaf segments were immersed in distilled water for 24 h, and finally dry weight (DW) was measured after leaf segments were dried at 65 °C in an oven for 24 h. Relative water content (RWC) was calculated as:

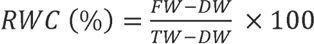

### Collection of tissue and chlorophyll measurements

Aboveground tissue from 8-12 plants were pooled together for each biological replicate and used for plastoglobule and thylakoid isolations, chlorophyll measurements, and proteomic and lipidomic analyses. Chlorophyll was measured spectrophotometrically according to Lichtenthaler (1987). Briefly, working in the dark under a green safety lamp at 4 °C, for each replicate 20 mg FW of leaf tissue was ground in liquid N_2_ and mixed with cold 80% (v/v) acetone. The resulting slurry was immediately transferred into a 1.5 mL Eppendorf tube. After incubating with gentle mixing overnight, the samples were centrifuged at 18,000 g for 15 minutes at 4 °C. The supernatant was diluted 10-fold in cold 80% (v/v) acetone and absorbance measured at 663.2, 646.8 and 470.0 nm wavelengths. Concentration estimates were calculated using the following equations:

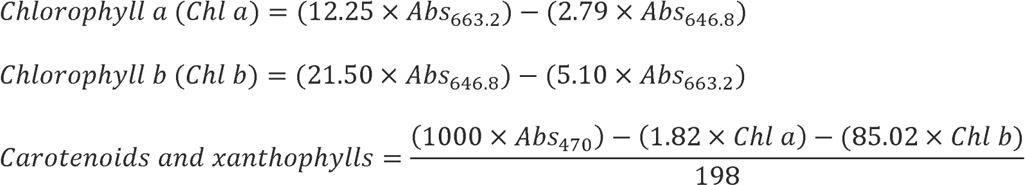

### Photosynthetic measurements

The fourth leaf was selected from eight randomly-selected plants at each time point for photosynthetic measurements. Measurements were collected from the middle of the leaf, off of the midrib, 2-3 hrs after lights turned on using the MultiSpeQ v2.0 device (PhotosynQ Inc., East Lansing, MI, USA) with the RIDES 2.0 parameters, available on the PhotosynQ platform (https://photosynq.org/). Data was plotted for photosynthetic activity (F_v_’/F_m_’), the total energy dissipation as non-photochemical quenching (NPQ_t_), and the quantum yield of photosystem II (F_II_). The photosynthetic measurements were uploaded to the PhotosynQ platform (https://photosynq.org/) with project ID: ‘DW-Peter-2021’ and are publicly available.

### Transmission electron microscopy

The fourth leaf of four randomly selected plants were trimmed at the midpoint of the leaf (halfway between base and tip and off of the midrib and sliced into 2 x 2 mm^2^ pieces with a fresh razor blade at the growth chamber before placing on ice for immediate delivery to the laboratory. Tissue sections were then fixed in 2% (w/v) paraformaldehyde and 2 % (w/v) glutaraldehyde in 0.1 M cacodylate buffer by infiltration under vacuum, and incubated overnight at room temperature. The fixed samples were washed in cacodylate buffer and post-fixed in 1% (w/v) osmium tetroxide in the same buffer for 2 h. They were then dehydrated in an ethanol series, transferred to propylene oxide, and embedded in Spurr’s resin. Thin sections (70Lnm) were cut with a diamond knife on a Power Tome ultramicrotome (RMC, Boeckler Instruments, Tucson, AZ, USA), and the sections were stained with 2% (w/v) uranyl acetate for 30Lmin and then with 3% (w/v) lead citrate for 15 min. Observations of chloroplasts were made on a JEM 1400Flash transmission electron microscope (JEOL, Tokyo, Japan) fitted with a high-resolution Tietz F224 digital camera. Morphological measurements from the micrographs were made using ImageJ software available from the web at http://rsbweb.nih.gov/ij/.

#### Plastoglobule and thylakoid preparations

Plastoglobules were isolated from the aboveground tissue by collecting 15 plants (around 80 g of tissue) for each biological replicate. Isolation was performed essentially as described in Shivaiah et al. (2022) was used for isolation of thylakoids and purification of plastoglobules. Briefly, the maize leaves were ground in a Waring blender in 500 mL of grinding buffer (50 mM HEPES-KOH (pH-8.0), 5 mM MgCl_2_, 100 mM Sorbitol) and filtered through gauze. Thylakoid membranes were pelleted by centrifugation of the filtrate at 1500 *g* for 6 min at 4 °C and were resuspended in medium R ([HEPES pH 8.0], 5 mM MgCl_2_, a cocktail of phosphatase and protease inhibitors containing antipain (74 mm) bestatin (130 mm), chymostatin (16.5 mm), E64 (56 mm), leupeptin (2.3 mm), phosphoramidon (37 mm), AEBSF (209 mm), aprotinin (0.5 mm), NaF (50 mM), b-glycerophosphate (25 mM), Na-orthovanadate (1 mM), and Na-pyrophosphate (10 mM)) with 0.2 M sucrose. Several aliquots of the resuspension were flash frozen in liquid N_2_ and stored at -80 °C until use. The remaining thylakoid membranes were aliquoted into four 3.2 mL polycarbonate open-top ultracentrifuge tubes (Beckman Coulter, Pasadena, CA, USA) and tip sonicated with a FB120 sonicator and a 4-tip horn (Fisher Scientific, Hampton, NH, USA). Each tube was pulse sonicated 4 x 10 sec at an amplitude (100%) with a 50 sec pause on ice between each pulse. Sonicated thylakoid suspensions were centrifuged at 150,000 x g for 30 min at 4 °C. The resulting floating pad consisting of crude plastoglobules were collected using a disposable syringe and 22G needle. Crude plastoglobules were then purified on a sucrose gradient of 0.7 M, 0.2 M and 0 M sucrose in medium R. Purified plastoglobules were collected from the top of the gradient, flash frozen, and stored at -80 °C until use.

### Lipid analysis (GC-FID) from thylakoids and PG’s

Lipids were extracted as described by Wewer et al. (2013). Briefly, thylakoid samples were adjusted to the equal protein concentration (∼0.1Lg) and homogenized in 5LmL of chloroform/methanol/formic acid (10:20:1, v/v/v); the homogenate was collected and shaken vigorously. Subsequently, 2.5 mL of 1 M KCl in 0.2LM H_3_PO_4_ was added and the mixture was vortexed briefly. The homogenized samples were centrifuged at 13000*g* for 1 min, and the lower chloroform layer was transferred to a new vial. Extraction was repeated three times by adding 5LmL of chloroform/methanol (2:1, v/v) to the residue, shaking and centrifuging the mixture, and collecting and pooling the chloroform phases. The combined chloroform phases were evaporated with a stream of N_2_ gas, and 100 μL of chloroform was added to dissolve the lipids. Polar lipids were separated with TLC plates treated with (NH4)_2_SO_4_ as described by Wang and Benning (2011). The galacto- and phospho-lipids were marked after iodine staining and scraped off with a razor blade and placed into 1.5 mL Eppendorf tubes. Using pentadecanoic acid (C15:0) as an internal standard, acyl groups were trans-methylated into their fatty acyl methyl esters (FAME) using 1 M HCl in methanol at 80 °C for 20 min. The resulting FAMEs were quantified by a gas-liquid chromatography system (GC-2010; Shimadzu Corp., Kyoto, Japan) equipped with a 30 m DB-23 column with inner diameter of 0.25 mm and film thickness of 0.25 µm (Agilent Technologies, Santa Clara, CA, USA) and with flame ionization detector (FID) according to Zhang et al. (2015).

### Immunoblots

Thylakoids and plastoglobules were separated by SDS-PAGE and solubilized according to Laemmli (Laemmli, 1970). Proteins were transferred on to nitrocellulose membrane using the semi-dry method and then probed with primary antibodies (Anti-PsbA, 1:10,000, anti-FBN1a, 1:5000, and anti-PsbS, 1:5000) purchased from Agrisera AB (product numbers: AS05 084, AS06 116 and AS03 032, respectively; Vännäs, Sweden) followed by incubation with secondary polyclonal anti-rabbit antibody tagged with HRP from Cell Signaling Technology (product number 7074; Danvers, MA, USA) and detected with chemiluminescence using the Amersham ECL kit (product number RPN2232; Chicago, IL, USA).

### Prenyl-lipid analysis

Thylakoids and plastoglobules were resuspended in 200 mL ice-cold methanol and mixed by inversion at 4 °C under dark for 5 mins. Then, 200 mL 50 mM Tris-HCl pH 7.5 and 1 M NaCl were added and incubated for an additional 10 min at 4 °C. For phase separation, 500 mL of ice-cold chloroform was added, and tubes were incubated for another 10 min at 4 °C. The mixture was centrifuged at 3000 g for 5 min at 4 °C. The hypophyses were collected with glass Pasteur pipette and transferred to a new tube. The extraction was repeated three times by adding ice-cold chloroform and mixture was incubated for 10 min and the phase separation done by centrifugation. The pooled chloroform was evaporated under gentle purging of N_2_ gas and stored under N_2_ gas in the dark at -20 °C until use. Immediately before use, prenyl lipids were resuspended in 350 mL of ethyl acetate.

Prenyl-lipids were analyzed as described previously in Espinoza-Corral *et al*. (2021). Briefly, samples were filtered through a 0.45-µm syringe filter and injected (20 µL) onto a reversed phase C30 chromatography column [AccuCore 2.6 µm beads (150 × 2.1 mm); Thermo Scientific, Waltham, MA, USA] using a Nexera-i LC-2040C 3D system (Shimadzu Corp., Kyoto, Japan) with a flow rate of 0.25 mL min^−1^ and a photodiode array detector (PDA) scanning continuously between 250 and 700 nm. A 30-min gradient program was used consisting of mobile phase A (methanol), mobile phase B (water), and mobile phase C (methyl tert-butyl ether) to separate prenyl-lipid compounds. Initial conditions were 80% A, 20% B, which was increased from 80% to 99% over 10 min, followed by a step to 94% A, 1% B and 5% C, a linear gradient to 34% A, 1% B, 65% C at 22 min, before returning to initial conditions by 27 min and an equilibration for 3 min. Peaks were identified by comparing against authentic standards and the absorbance spectrum. Peak identities were determined within the Lab Solutions software package (version 5.97 SP1, Shimadzu Corp., Kyoto, Japan) by comparing their retention time and absorbance spectrum with calibration curves derived from commercial standards, except for plastoquinone (PQ). The quantification of PQ was estimated using the extinction coefficient and calibration curve developed from the ubiquinone-10 commercial standard.

### LC-MS/MS lipid measurements

Total lipids were extracted from freshly isolated thylakoids (5 mg of protein) and freshly isolated plastoglobules (OD_700_ = 1.0) with 1 mL of extraction buffer (chloroform:methanol:formic acid (20:10:1 v/v/v)). Samples were shaken vigorously for 5 min, and then 500 mL of water: methanol (3:1 v/v) was added and well mixed. Phases were completely separated by centrifugation for 2 min at 13 000 *g* at room temperature. The upper chloroform phase was removed and lipids re-extracted three times. Pooled chloroform phases were dried down under a gentle stream of N_2_ gas. Lipids were resuspended in 100 µL isopropanol and diluted 20-fold for LC-MS/MS measurement.

Lipid samples were analyzed using a Waters Xevo G2-XS Q-TOF mass spectrometer interfaced with a Waters Acquity binary solvent manager and Waters 2777c autosampler. Next, 10 µL samples were injected onto an Acquity UPLC BEH-C18 column (2.1 x 100 mm) held at 55 °C. The mobile phases consisted of 0.1% (w/v) formic acid and 10 mM ammonium formate in acetonitrile/water (60:40 v/v) (Solvent A) and isopropanol/acetonitrile (90:10, v/v) containing 0.1% (w/v) formic acid and 10 mM ammonium formate (Solvent B). A gradient of mobile phase was applied in a 20-min program with a flow rate of 0.4 mL min ^-^1. The gradient profile was performed as follows: initial conditions were 80% A and 20 % B, ramp to 43% B at 2 min , followed by a linear ramp to 54% B at 12 min, ramp to 70% B at 12.10 min, ramp to 99% B at 18 min, then return to 20% B at 18.1 min and hold until 20 min. LC separated analytes were ionized by positive ion mode electrospray ionization and mass spectra were acquired using an MSe method in continuum mode over m/z 50 to 1500 to provide data under non-fragmenting and fragmenting conditions (collision energy ramp from 20–80 V). Capillary voltages were set to 3 kV (positive mode) or 2 kV (negative mode), cone voltage was 30 V, source temperature was 100°C, desolvation temperature was 350°C, desolvation gas flow was 600 L/hr and cone gas flow was 40 L/hr.

### Proteomics sample preparation

Proteomic analysis was performed with lyophilized thylakoids normalized to an equal amount of protein (250 µg), lyophilized plastoglobules normalized to 1.0 OD_700_ or homogenized total leaf samples normalized to equal protein concentration (200 µg). Total leaf, thylakoids and plastoglobules were re-suspended in Laemmli sample buffer and heated at 60 °C for 10 min.

Samples were cooled and loaded onto a 12.5% pre-cast Criterion 1D gel (Bio-Rad, Hercules, CA, USA) and electrophoresed at 50 V until the Coomassie dye front migrated 2-3 mm below the well. The gel was stained using Coomassie Brilliant Blue G250 (0.05% (w/v) in 50% (v/v) methanol, 10% (v/v) glacial acetic acid and 40% (v/v) water. Concentrated sample bands were then excised from the gel and placed into individual microfuge tubes.

Gel bands were digested in-gel according to Shevchenko, et al. (1996) with modifications. Briefly, gel bands were dehydrated using 100% acetonitrile and incubated with 10 mM dithiothreitol in 100 mM ammonium bicarbonate, pH∼8, at 56°C for 45 min, dehydrated again and incubated in the dark with 50 mM chloroacetamide in 100 mM ammonium bicarbonate for 20 min. Gel bands were then washed with ammonium bicarbonate and dehydrated again. Sequencing-grade modified trypsin was prepared to 0.005 µg/µL in 50 mM ammonium bicarbonate and ca. 100 µL of this was added to each gel band so that the gel was completely submerged. Bands were then incubated at 37 °C overnight. Peptides were extracted from the gel by water bath sonication in a solution of 60 % (v/v) acetonitrile (ACN) / 1% (v/v) trifluoroacetic acid (TFA) and vacuum dried to ca. 2 µL.

### LC-MS/MS proteomics

The peptides were resuspended in 20 µL of a solution of 2% (v/v) acetonitrile, 0.1% (v/v) triflouroacetic acid and 97.9% (v/v) water. A sample injection of 5 mL was made using a Thermo (www.thermo.com) EASYnLC 1200 onto a Thermo Acclaim PepMap RSLC 0.075mm x 250mm C18 column and washed for ca. 5 min with buffer A (99.9% water / 0.1% formic acid; v/v). For total lead and thylakoid samples, bound peptides were then eluted over 45 min with a linear gradient of 8% buffer B (80% acetonitrile / 0.1% formic acid / 19.9% water; v/v/v) to 35% buffer B in 34 min at a constant flow rate of 300 nL min^-1^. After the gradient the column was washed with 90% buffer B for the duration of the run. For plastoglobule samples, bound peptides were eluted over 35 min with a linear gradient of 8% buffer B to 35% buffer B in 24 min at a constant flow rate of 300 nL min^-1^. After the gradient the column was washed at 90% buffer B for the duration of the run. Column temperature was maintained at a constant temperature of 50 °C using and integrated column oven (PRSO-V2, Sonation GmbH, Biberach, Germany). Eluted peptides were sprayed into a ThermoScientific Q-Exactive HF-X mass spectrometer (www.thermo.com) using a FlexSpray spray ion source. Survey scans were taken in the Orbitrap (60000 resolution, determined at m/z 200) and the top 15 ions in each survey scan were subjected to automatic higher energy collision induced dissociation with fragment spectra acquired at a resolution of 15000.

### LC-MS proteomic data analysis

The resulting MS/MS spectra were converted to peak lists using MaxQuant v1.5.5.1 (Cox and Mann, 2008) and searched against the *Z. mays* (v3) full peptide dataset (63241 unique protein sequences downloaded from MaizeGDB [https://www.maizegdb.org/]), and concatenated with common laboratory contaminants using the Andromeda search algorithm (Cox et al., 2011). Oxidation of methionine, phosphorylation of threonine, serine, and tryptophan, and N-terminal acetylation were set as variable modifications, carbamidomethylation was set as a fixed modification. Digestion mode was Trypsin/P with a maximum of two missed cleavages. Label-free quantification was enabled with default settings. MS/MS tolerance of the first search was 20 ppm, and the main search was 4.5 ppm, with individualized peptide mass tolerance selected. FDR at peptide spectrum match and protein levels was set as 0.01. Total leaf, plastoglobule and thylakoid samples were independently searched in separate groups of 0.7 min match time window and 20 min alignment window for match between runs. The ‘primary gene model’ for each protein group was identified as that with the highest number of Unique + Razor Peptide counts when compiled across all experiments. In the case of a tie, the lowest gene model number was selected, regardless of whether the protein group contained gene models from one or more gene loci. The mass spectrometry proteomics data have been deposited to the ProteomeXchange Consortium via the PRIDE (Perez-Riverol et al., 2022) partner repository (https://www.ebi.ac.uk/pride/) in MIAPE-compliant format with the dataset identifiers PXD050599 and PXD057023.

### Graphing and statistical analyses

Statistical analyses and plotting were performed with Prism version 9.3 (GraphPad, Boston, MA, USA). Data on subjected to statistical analysis by Minitab (Minitab, State College, PA, USA) with one- way ANOVA or Student’s t-test.

## ACKNOWLEDGEMENTS & FUNDING

This work is supported by the Plant Health and Production and Plant Products program, project award no. MICL08607, from the U.S. Department of Agriculture’s National Institute of Food and Agriculture to P.K.L.. We thank Drs. Sheng Ying and Kiran-Kumar Shivaiah for assistance with data collection, Dr. Qianjie Wang for assistance with the MaxQuant analysis, Doug Whitten of the MSU Proteomics Core Facility for assistance with proteomics studies, and Alicia Withrow of the MSU Center for Advanced Microscopy for assistance with collection and imaging of transmission electron micrographs.

## SUPPLEMENTARY MATERIALS

**Supplementary Figure S1**. Visualization of whole plant (**A**) and leaf (**B**) phenotypes during well-watered control treatment and water-deprivation treatment.

**Supplementary Figure S2**. Measurements of chloroplast and plastoglobule ultrastructure from M cells (left column) and BS cells (right column) from transmission electron micrographs.

**Supplementary Figure S3**. Isolation and enrichment of plastoglobules and thylakoids.

**Supplementary Figure S4**. Summed nLFQ intensities of all detected subunits of specified photosynthetic complexes or protein families in thylakoid samples.

**Supplementary Figure S5**. nLFQ intensities of maize Fibrillin 4 (FBN4) and the sum of the Fibrillin 7 paralogs (ΣFBN7) in plastoglobules (left) and thylakoids (right).

**Supplementary Table S1.** Leaf relative water content at each time point

**Supplementary Table S2.** Pigment measurements at each time point

**Supplementary Table S3.** Photosynthetic measurements at each time point

**Supplementary Table S4.** Total leaf proteomics analysis

**Supplementary Table S5.** Cellular ultrastructure measurements

**Supplementary Table S6.** All Protein Identifications from Isolated Thylakoid (with associated plastoglobules)

**Supplementary Table S7.** All protein identifications from isolated plastoglobules

**Supplementary Table S8.** Quantitative plastoglobule proteome at each time point

**Supplementary Table S9.** Polar lipid ratios of isolated plastoglobules

**Supplementary Table S10**. Acyl chain composition of polar lipids

**Supplementary Table S11.** Thylakoid LC-MS/MS lipidome of *Z. mays* leaf tissue

**Supplementary Table S12.** Plastoglobule LC-MS/MS lipidome from *Z. mays* leaf tissue

