## Supplementary Figures for "Dynamic changes to the plastoglobule lipidome and proteome in water-deficient maize"

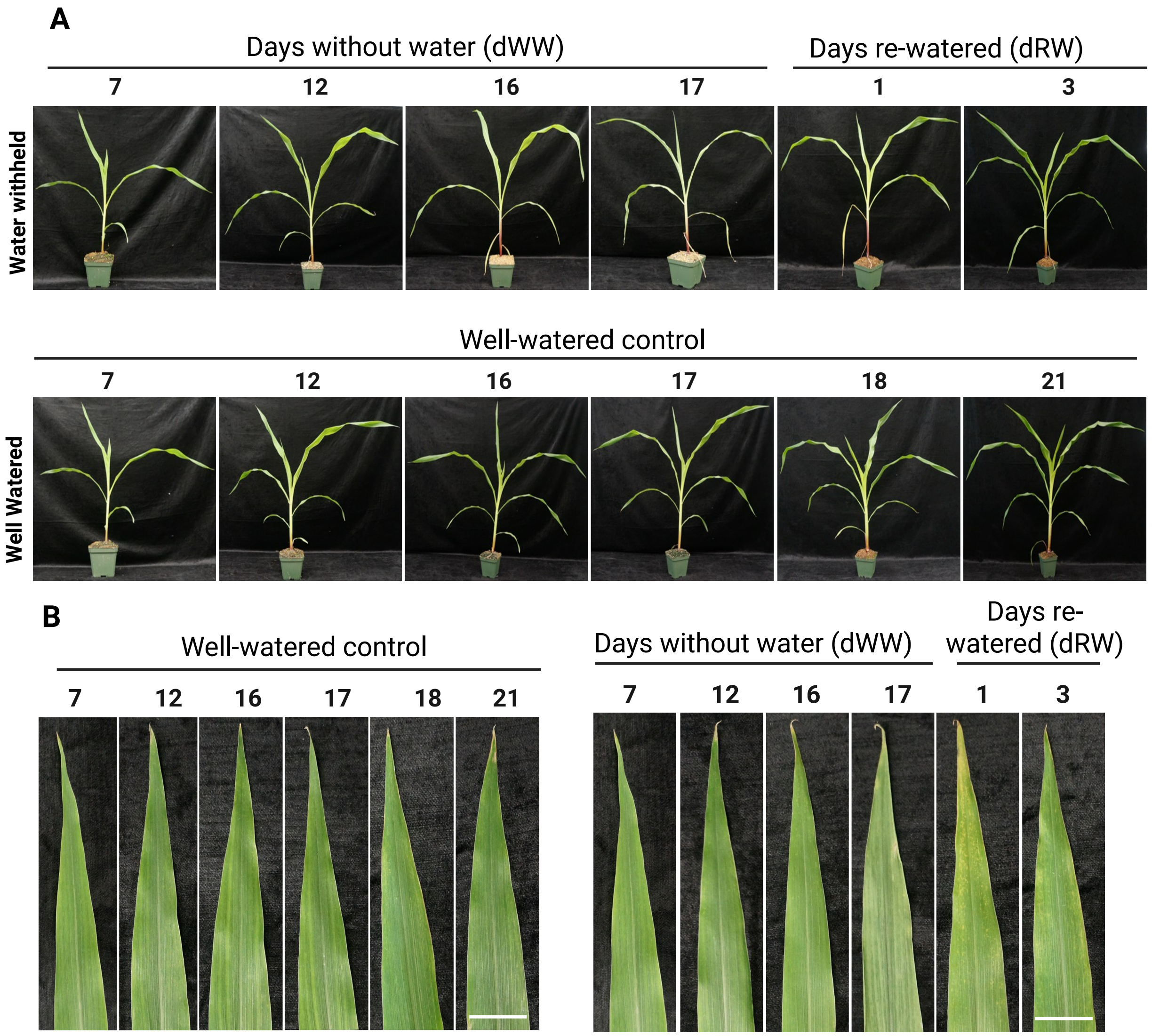


**Supplementary Figure S1**. Visualization of whole plant (**A**) and leaf (**B**) phenotypes during well-watered control treatment and water-deprivation treatment. The fourth leaf was selected from each plant and the white scale bar indicates 0.5 cm.

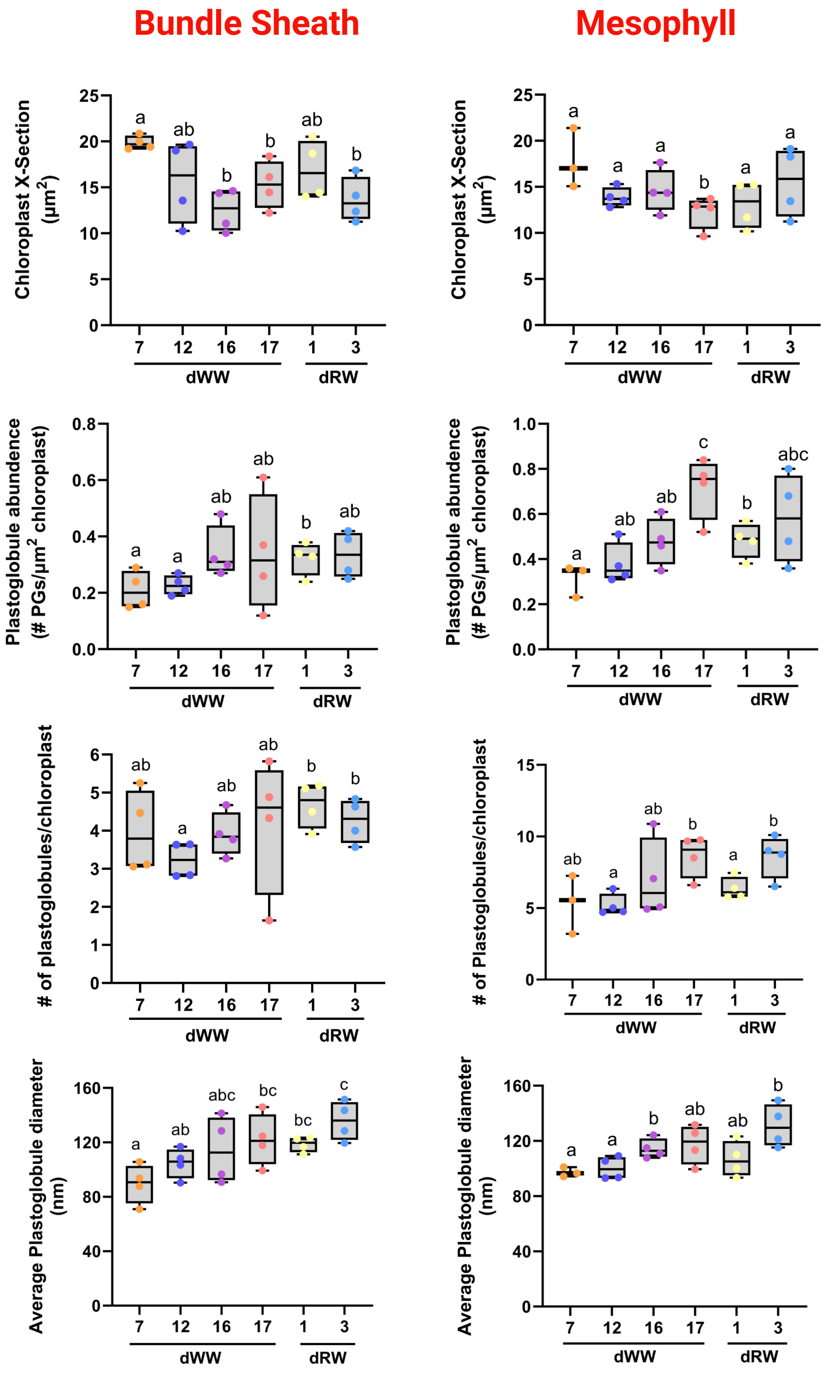


**Supplementary Figure S2**. Measurements of chloroplast and plastoglobule ultrastructure from M cells (left column) and BS cells (right column) from transmission electron micrographs. All measurements are plotted as box and whisker plots in which the box limits indicate 25th and 75th percentiles, horizontal line is the median, and whiskers display minimum and maximum values and overlaid with individual data points. Micrographs were analyzed manually in Image. Statistically significant differences were identified by homoscedastic two-tailed Student's t-test ($p < 0.05$), and are indicated by lower case letters, n = 4 biological replicates, *i.e.* individual plants (each data point representing the average values from a minimum of two micrographs from an individual plant).

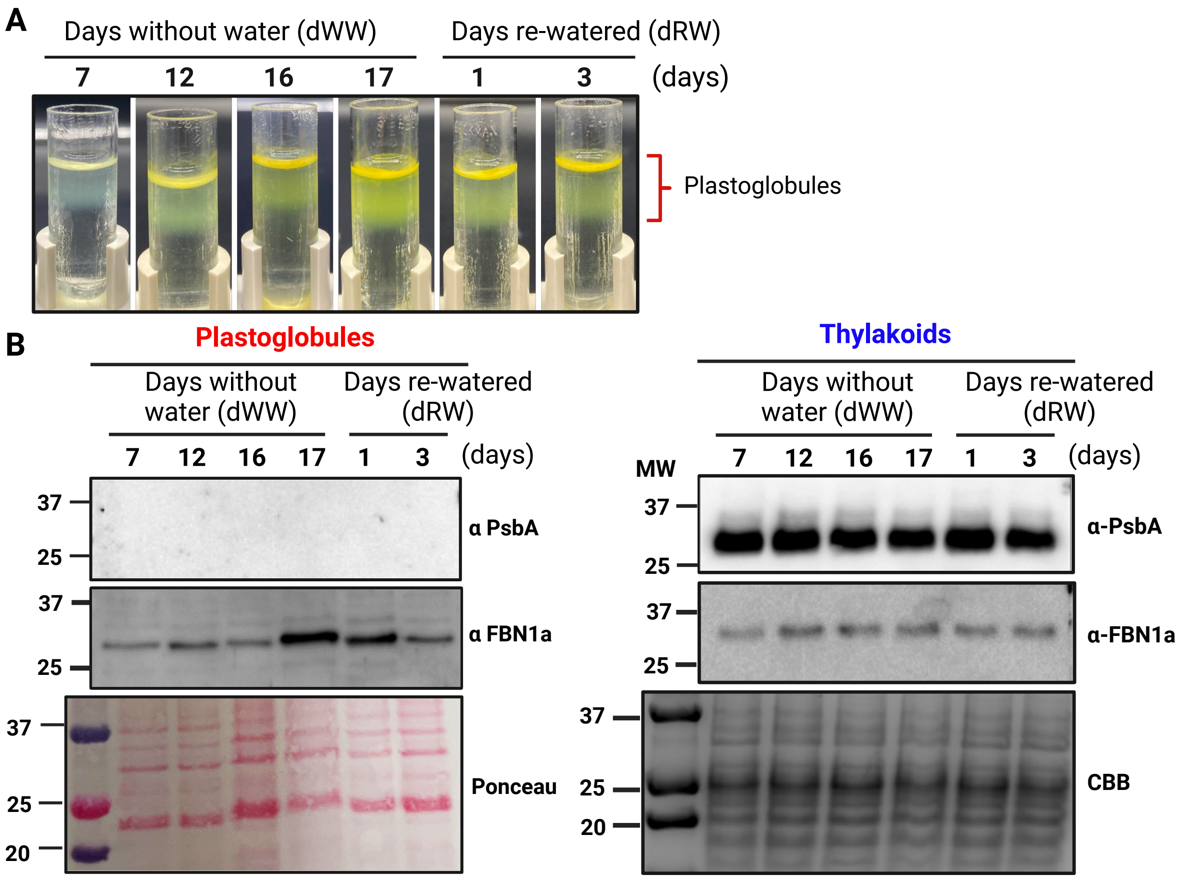


**Supplementary Figure S3**. Isolation and enrichment of plastoglobules and thylakoids. **A)** Representative images of floating plastoglobule isolations following ultracentrifugation on a sucrose gradient. Plastoglobules are seen as the yellow floating pads indicated with the red bracket on the right side. This materials was subsequently harvested from the gradient and saved as purified plastoglobule samples. **B)** Immunoblots of isolated plastoglouble (left) and thylakoid (right) demonstrating the enrichment and purity of the samples.

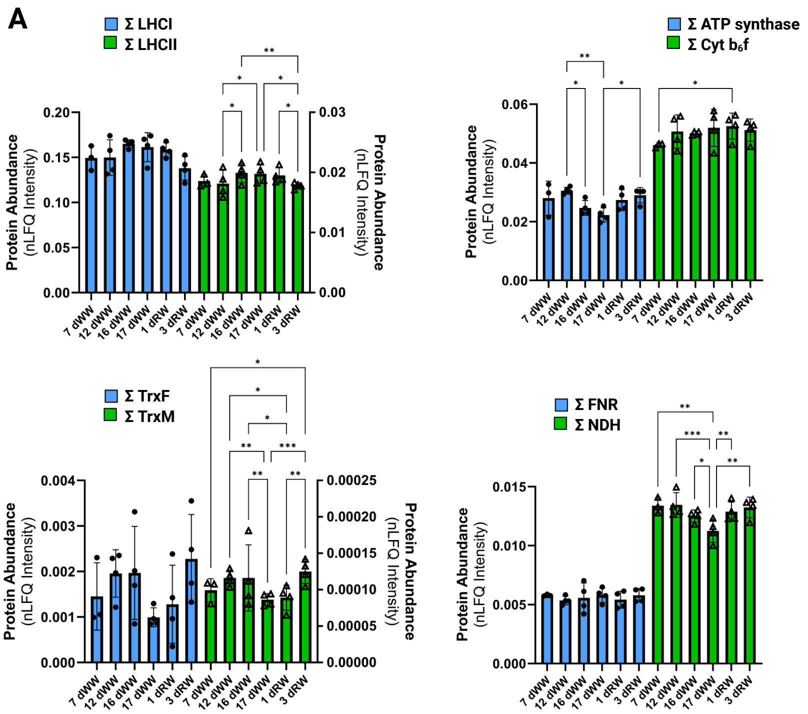


**Supplementary Figure S4**. Summed nLFQ intensities of all detected subunits of specified photosynthetic complexes or protein families in thylakoid samples. Graphs plot mean ± 1 standard error of the mean, n = 3 (7 dWW) or 4 (all other time points) biological replicates (*i.e.*, individual plants). Statistically significant differences were determined using a mixed effects model (panel A) or ordinary two-way ANOVA (panel B); * $p < 0.033$, ** $p < 0.002$, *** $p < 0.001$.

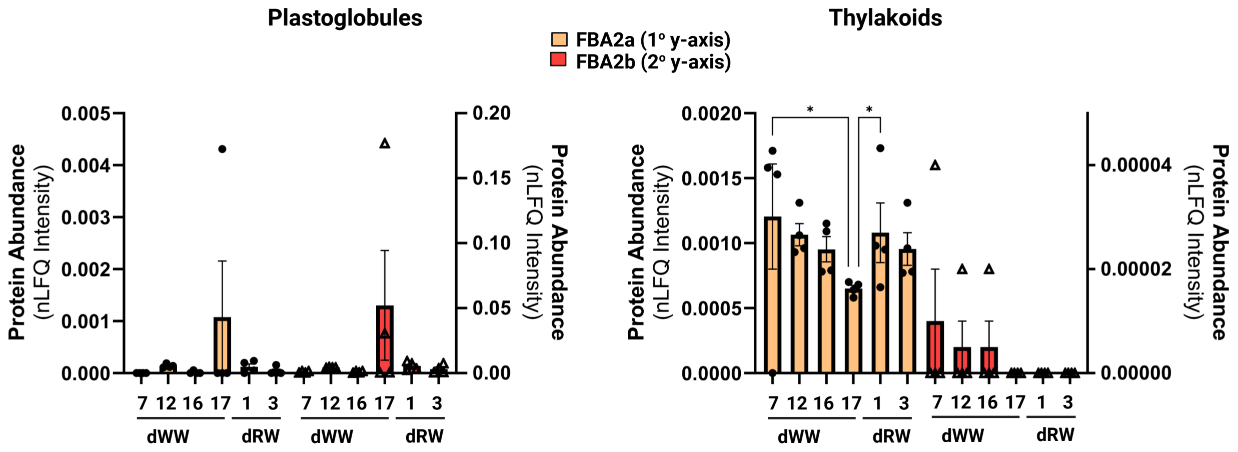


**Supplementary Figure S5**. nLFQ intensities of maize Fructose bis-phosphate Aldolase isoforms 2a and 2b (FBA2a/sb) in plastoglobules (left) and thylakoids (right). Graphs plot mean ± 1 standard error of the mean, n = 3 (7 dWW) or 4 (all other time points) biological replicates (*i.e.*, individual plants). FBA2b is plotted on the secondary y-axis. Statistically significant differences within protein groups were determined using an ordinary two-way ANOVA; * $p < 0.033$, ** $p < 0.002$, *** $p < 0.001$.

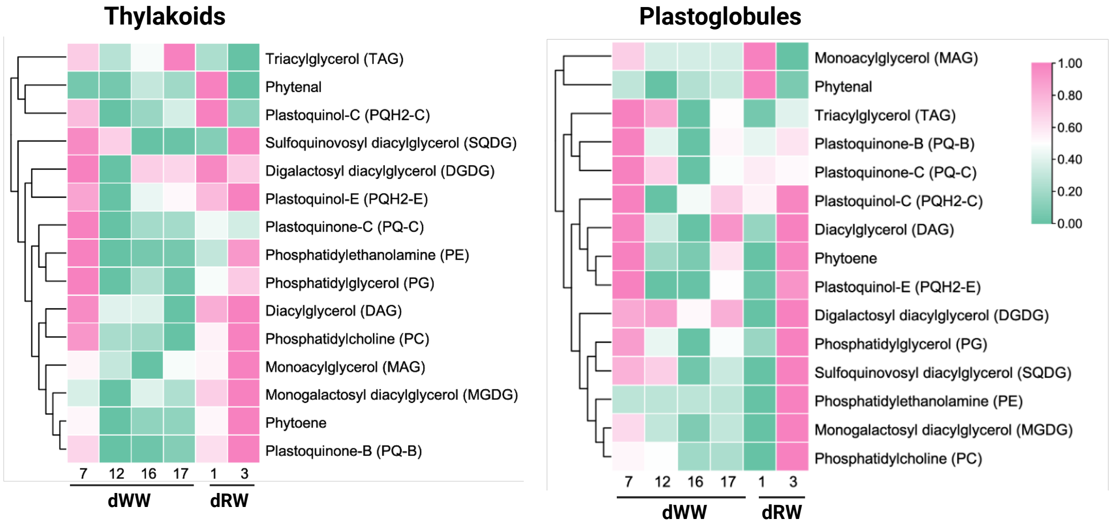


**Supplementary Figure S6.** Relative levels of polar-, neutral-, and prenyl-lipids in *Z. mays* plastoglobules and thylakoids. Mean values plotted, n = 3 biological replicates (*i.e.*, individual plants). TAG, triacylglycerol ; DAG, diacylglycerol; MAG, monoacylglycerol. Heat map representing the $\log_2$ values of each of the annotated triacylglycerol species in thylakoids (left) and plastoglobules (right) at each time point. For more details about the data, refer to Supplementary Tables S11 and S12.
